# Cryo-EM Heterogeneity Analysis using Regularized Covariance Estimation and Kernel Regression

**DOI:** 10.1101/2023.10.28.564422

**Authors:** Marc Aurèle Gilles, Amit Singer

## Abstract

Proteins and the complexes they form are central to nearly all cellular processes. Their flexibility, expressed through a continuum of states, provides a window into their biological functions. Cryogenic electron microscopy (cryo-EM) is an ideal tool to study these dynamic states as it captures specimens in non-crystalline conditions and enables high-resolution reconstructions. However, analyzing the heterogeneous distributions of conformations from cryo-EM data is challenging. We present RECOVAR, a method for analyzing these distributions based on principal component analysis (PCA) computed using a REgularized COVARiance estimator. RECOVAR is fast, robust, interpretable, expressive, and competitive with the state-of-art neural network methods on heterogeneous cryo-EM datasets. The regularized covariance method efficiently computes a large number of high-resolution principal components that can encode rich heterogeneous distributions of conformations and does so robustly thanks to an automatic regularization scheme. The novel reconstruction method based on adaptive kernel regression resolves conformational states to a higher resolution than all other tested methods on extensive independent benchmarks while remaining highly interpretable. Additionally, we exploit favorable properties of the PCA embedding to estimate the conformational density accurately. This density allows for better interpretability of the latent space by identifying stable states and low free-energy motions. Finally, we present a scheme to navigate the high-dimensional latent space by automatically identifying these low free-energy trajectories. We make the code freely available at https://github.com/ma-gilles/recovar.

---

Proteins and their complexes play pivotal roles in cellular processes, governing essential biological functions. The function of these biological macromolecules can be elucidated through the continuum of their structural states encompassing local dynamics, large-scale rearrangement of domains, and modification of subunits.

Cryogenic electron microscopy (cryo-EM) has emerged as a powerful tool for investigating the dynamic conformational landscapes of biomolecules, recognized as one of Nature Methods’ “Methods to Watch” in 2022 [10]. By directly imaging biological specimens in a near-native, frozen-hydrated state, cryo-EM circumvents the need for crystallization, a process that can distort molecular structure and restrict conformational variability. This unique approach allows cryo-EM to capture a more accurate representation of biomolecules in their functional states, revealing the full spectrum of their conformational ensemble. Moreover, cryo-EM’s ability to generate high-resolution images of individual particles enables the identification of subtle conformational differences within a population, providing unprecedented insights into biomolecular dynamics. Cryo-EM can tackle large and complex assemblies, making it a preferred method for exploring conformational heterogeneity.

However, the computational and modeling challenges posed by deciphering the heterogeneous distribution of conformations from cryo-EM datasets are considerable. The workhorse of cryo-EM heterogeneity analysis is 3D-classification [44], which aims to discern discrete conformational states. While effective for a few evenly distributed states, this approach encounters difficulties with many states or uneven distributions of discrete states and cannot capture continuous changes.

A diverse array of methods has been proposed to address these challenges. These methods encompass rigid body fitting [32], linear subspace methods [55, 23, 35, 2, 38], deep-learning approaches [65, 4], manifold learning methods [13, 31, 48], strategies based on molecular dynamics [59], and methods focused on computing deformation maps [19, 39, 47]. For a comprehensive review of these rapidly evolving methodologies, refer to [53, 58, 11]. Though diverse in their details, most of these methods share a common framework: they employ a mapping to embed images into a “latent space”. This mapping aims to isolate image variations stemming from irrelevant factors like pose, imaging effects, and noise, focusing solely on variability due to conformational changes. A second mapping then translates this latent space into conformational states.

Significant advancements have been made recently, particularly with the utilization of neural networks producing volumes at high resolution. However, present methods grapple with several issues, including a lack of explainability and interpretability of the latent space, the absence of regularization, and numerous hyperparameters requiring tuning. These challenges are further exacerbated by the absence of metrics to validate or compare results from cryo-EM heterogeneity analysis; particularly for some deep-learning methods that have shown the ability to hallucinate physically plausible motion (see fig. A.11). Consequently, distinguishing between a genuinely high-resolution reconstruction and a mere overfit artifact remains difficult. Additionally, the reconstruction phase is often only the beginning of the analysis. Estimating the free energy of states or motions is often the ultimate goal. Both quantities are typically calculated based on recovering the likelihood of observing each state in the data, usually numerically estimated by density estimation in latent space. However, most methods for generating latent space do not conserve distances or densities. Thus, the density estimates obtained this way can be entirely unrelated to the data and instead be artifacts of the method [26, 58].

To tackle these challenges, we introduce RECOVAR, a method grounded in principal component analysis (PCA) computed via REgularized COVARiance estimation. This method is white-box, automatically regularized, allows for the estimation of density, and offers results that outperform neural networks in terms of resolution. We overcome overfitting and parameter-tuning issues using a Bayesian framework, building upon successful schemes for homogeneous reconstructions [45]. Our method belongs to the family of linear subspace methods, which have achieved practical success thanks to 3DVA [38], implemented in the popular software suite cryoSPARC [40]. However, linear subspace methods have often been considered limited to low resolution [54] or suitable only for capturing “small” motion and “simple” heterogeneity. We demonstrate that this perception is primarily based on misunderstanding their properties. Linear subspace methods do not capture the inherent heterogeneous dimension but are more akin to a change of basis. Specifically, PCA computes the optimal truncated basis to represent the heterogeneity and expresses conformations by their projections in this coordinate system. As a result, a large one-dimensional motion is typically not well-embedded in a one-dimensional subspace. However, we show that simply computing larger subspaces overcomes this limitation. Furthermore, we overcome the low-resolution concerns of PCA methods by introducing a novel scheme for reconstructing volumes based on an adaptive kernel regression, which aims to optimally combine the individual contribution of images across the frequency and spatial domain. Independent and extensive benchmarking of seven fixed-pose heterogeneity methods have found that this new volume generation scheme outperforms neural network and classical methods across nearly all signal-to-noise ratio (SNR), type of heterogeneity, and complexity of heterogeneity (RECOVAR attains the highest resolution scores on 11 out of 12 reported scores, see [21] for detailed analysis).

Additionally, we show how to leverage the favorable properties of linear subspace methods and the statistical framework to accurately estimate the density of the conformational distribution from the PCA embedding. Using this accurate density estimation method, we identify stable states and low free-energy motions from their high density in several real and synthetic datasets. Finally, we design a method to automatically identify high-density trajectories to recover these low free energy motions from high-dimensional latent spaces.

## 1 Results

### 1.1 Regularized covariance estimation

The method presented here uses PCA to compute an optimal linear subspace to represent the heterogeneous distribution of states. However, applying PCA to conformational states in cryo-EM data introduces a significant complexity: the observations are projection images of the conformational states, not the states themselves. Therefore, more than a straightforward application of PCA is needed. The principal components are also the eigenvectors of the covariance matrix of conformations, and remarkably, this covariance matrix can be estimated directly from projection images [23, 2]. Hence, as depicted in fig. 1^1^, our pipeline initiates with the statistical estimation of the mean and covariance of the conformational states directly from projection images. These estimation problems are regularized by splitting the data into halfsets and extending the Fourier Shell Correlation (FSC) regularization scheme ubiquitously used in homogeneous reconstruction [45]. Once computed, an eigenvalue decomposition of the covariance matrix provides the principal components. In the subsequent stage, we employ a Bayesian framework to infer each image’s probable conformation distribution within the principal components. Using the favorable properties of this embedding, we generate high-resolution volumes by careful image averaging, estimate the conformational density, and compute low free-energy trajectories.

**Figure 1.**
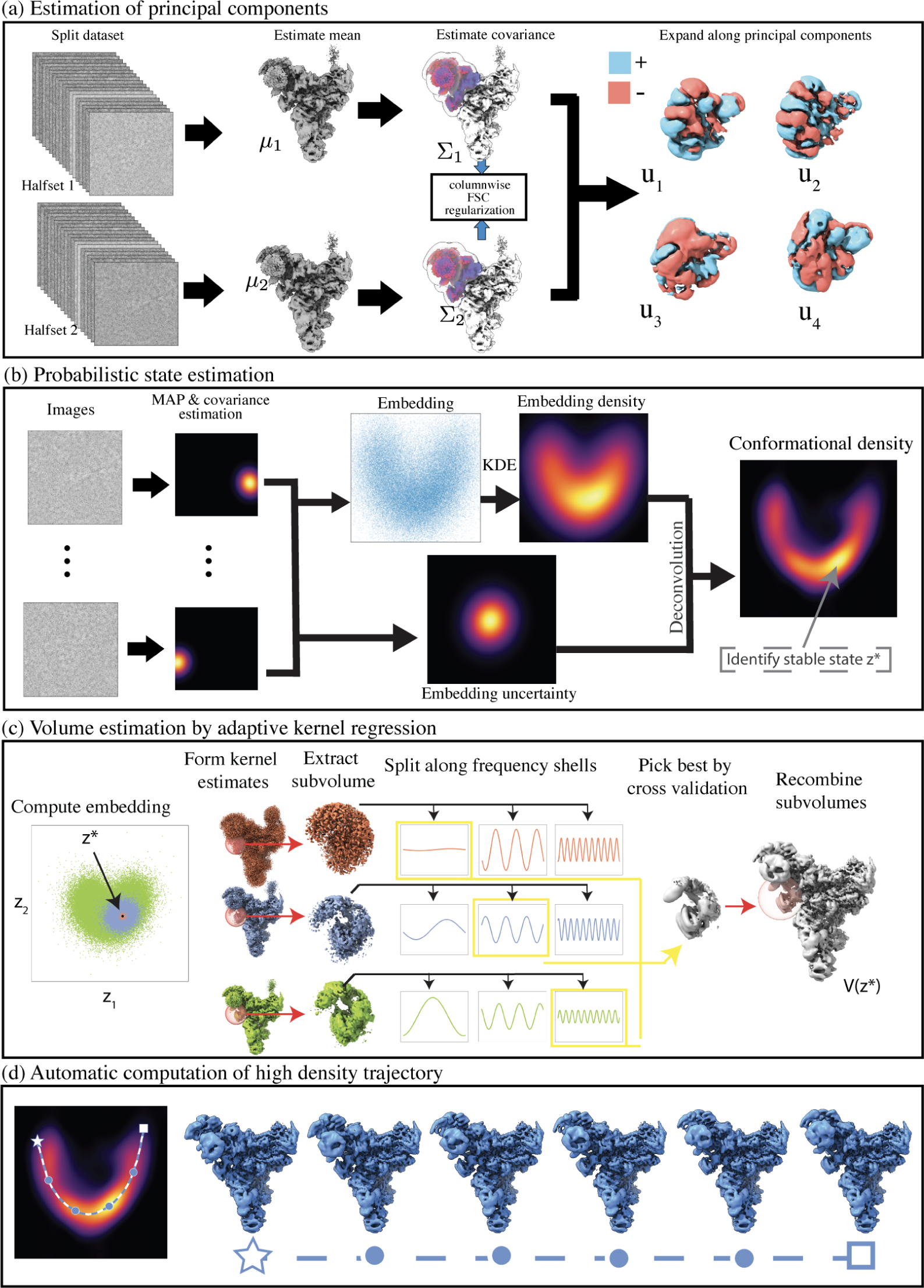
Illustration of the RECOVAR pipeline for heterogeneity analysis in cryo-EM on the snRNP spliceosome complex (EMPIAR-10073 [33]). **(a)** The covariance of the distribution of states is estimated and regularized using the FSC between two reconstructions from two halfsets. The principal components are then computed. **(b)** Each image is assigned a probabilistic label using maximum a posteriori (MAP) estimation. The uncertainty and the distribution of the labels, estimated by kernel density estimation (KDE), are used to infer the conformational density. The conformational density reveals the stable (high-density) conformational states. **(c)** Volumes are reconstructed from the embedding by adaptive kernel regression. **(d)** Motions are generated by computing high-density trajectories between an initial and final state (indicated by a star and a square). Intermediate reconstructed volumes along the trajectory are displayed.

The image formation process of cryo-EM expressed in the Fourier domain relating the Fourier transform of the 2-D image 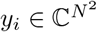 to the Fourier transform of the 3-D conformation 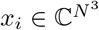, is typically modeled as:

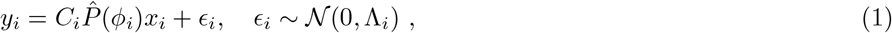

where 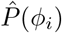 models the tomographic projection from 3-D to 2-D after rigid body motion parametrized by *ϕ*_*i*_, *C*_*i*_ models the contrast transfer function (CTF), and *ϵ*_*i*_ is Gaussian noise with covariance Λ_*i*_. In typical cryo-EM reconstruction, the poses *ϕ*_*i*_ are unknown and need to be inferred, often through a scheme alternating between inferring states *x*_*i*_ and poses *ϕ*_*i*_. In what follows, we assume that poses *ϕ*_*i*_ have been previously estimated, typically from a consensus reconstruction as is a common assumption for heterogeneity methods [65, 38], and fix the linear maps 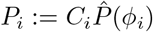, which model both CTF and projections. This assumption is restrictive as the poses cannot be estimated from consensus reconstructions for some highly heterogeneous cryo-EM datasets. For datasets where poses can be estimated from consensus reconstructions, alignment will likely not be perfect, and the residual pose errors can result in lower-than-possible resolution or even spurious heterogeneity. While some neural methods [27, 66] have shown the ability to infer poses ab-initio on some datasets, robust pose estimation for highly heterogeneous datasets remains an open problem which we come back to in the discussion.

When poses are known, the mean 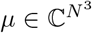 of the distribution of conformations can be straightforwardly estimated, e.g. by solving a linear least-squares problem:

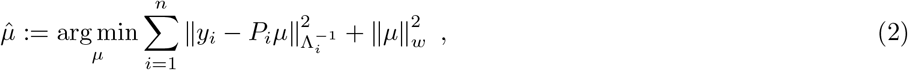

where 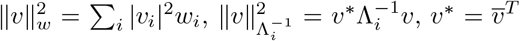 denotes the conjugate transpose, 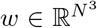 are optional Wiener filter parameters, *n* is the number of images and 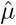 is the estimate of the mean conformation. Equation (2) is also the problem usually solved during homogeneous reconstruction. In that case, the weights *w* are often set using the FSC regularization scheme [45]; for example, it is the default method in popular cryo-EM software such as RELION [45] and cryoSPARC [40].

Analogously, the covariance of conformations can be estimated as the solution to the linear system corresponding to the following least-squares formulation [2, 23]:

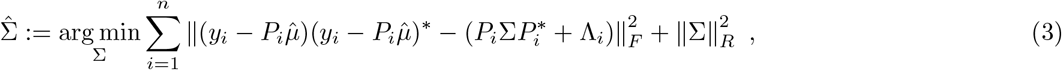

where 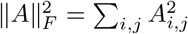 and 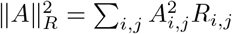 and *R* are the regularization weights. Unfortunately, the computation of this covariance estimator is a massive computational burden; e.g., representing conformations on a grid of size 128^3^ would result in a covariance matrix with 128^6^ entries—or 17 terabytes in single-precision floating-point. One solution to this problem is to use probabilistic PCA (PPCA) [57, 55] or similar heuristics [38] to compute a different linear subspace computed by an iterative procedure, see appendix A.9 for a discussion of the difference of these approaches.

We take a different route that does not require iteration—an expensive computation for large cryo-EM datasets—and instead computes the principal component using modern linear algebra techniques. First, we estimate only a subset of the entries of the covariance matrix and compute them using kernel regression. Then we use the fact that, for low-rank covariance matrices, only a subset of the columns is required to estimate the entire matrix and its leading eigenvectors, which are the principal components we seek. This celebrated fact in computational mathematics is the basis of numerical schemes such as the Nyström extension [61], see appendix A.2.

Entries of 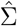 are estimated with a high dynamic range of SNR. This is analogous to the homogeneous reconstruction problem: low-frequency coefficients are easier to estimate as they are more often observed in images due to the Fourier slice theorem, and the SNR is higher at low frequencies. The covariance estimation problem presents an even wider range of SNR across pairs of frequencies [23]; making careful regularization even more crucial. To that end, we generalize the FSC regularization used in homogeneous reconstruction to the covariance estimation problem by carefully accounting for the highly nonuniform sampling of entries of the covariance matrix. This regularization proceeds by splitting the dataset into two halves and computing two independent copies of the same object. The correlation scores between the two copies provide estimates of the SNR, which can be used to set the regularization weights; see appendix A.3 for details. Optionally, we use a real-space mask to focus the analysis on one part of the molecule.

These computational advances, coupled with new regularization strategies for the covariance matrix, allow us to robustly and efficiently compute even a large number of principal components of principal components at high resolution that can encode a rich distribution of heterogeneous conformations. For example: for a dataset of 300,000 images of size 256^2^, on one GPU^2^, RECOVAR computes 100 principal components in 4 hours, where 3DVA takes 16 hours to compute only 20 principal components, and cryoDRGN takes 23 hours to train a network. This alleviates a limitation of linear subspace methods: their sometimes limited representation power for low-dimensional subspaces or low-resolution principal components.

### 1.2 Latent space embedding

We next turn to the problem of estimating the conformation present in each image. Due to noise and projection ambiguity, confidently matching an image to a single conformation is often impossible. Instead, we use a Bayesian framework to find a distribution of likely states for each image; Under the likelihood model in eq. (1), the posterior probability that an image comes from a particular state can be computed using Bayes’ law:

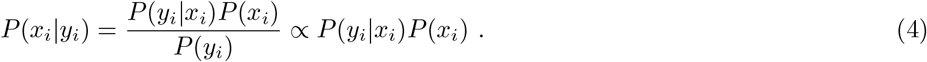

We approximate the distribution of states as a normal distribution 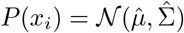, plug the truncated eigenvalue expansion of the estimated covariance matrix 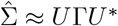 where the columns of 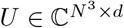 are the estimated principal components and the diagonal entries of Γ are the estimated eigenvalues, and parametrize *x*_*i*_ by its coordinates in the low-dimensional basis 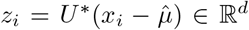, where *d* is the number of chosen principal components. We refer to the coordinate *z*_*i*_ as living in latent space and refer to states 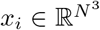 as living in volume space to make the distinction. We can now estimate explicitly a distribution of possible states *z*_*i*_ present in a given image *y*_*i*_:

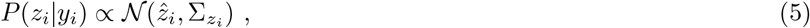

where 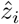 and 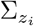 are the mean and covariance estimates of the conformation in image *i*, with explicit formulas given in appendix A.5. The covariance conformation estimate 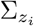 reflects the uncertainty in the latent variable assigned to image *i*, and is leveraged to generate volumes and to estimate the conformational density from the embedding^3^, see sections sections 1.3 and 1.4.

We further allow for contrast variation in each image by computing a per-image optimal scaling parameter similar to [38, 50], and describe a method to estimate this parameter at virtually no extra cost; see appendices A.4 and A.5 for details, and fig. A.7 for an illustration of the effect of this correction of downstream tasks.

### 1.3 Reconstructing conformations from embeddings

After the dataset is embedded into a common coordinate system, heterogeneity methods use a wide range of mappings to translate this latent space into volumes that can be visualized and interpreted. We highlight two choices of mappings here.

One approach used with linear embeddings, which we name *reprojection*, employs the explicit inverse map 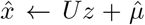, which entails taking a linear combination of principal components. This reprojection scheme only works well when the SNR is sufficiently high to compute all the relevant principal components. ^4^ In practice, limited SNR often allows access to only a subset of necessary principal components, and it is preferable to generate volumes by a different method; here we present a scheme based on adaptive kernel regression that averages the contribution of images by carefully weighting individual frequencies of each image; generalizing successful schemes for homogeneous reconstruction.

To resolve the high-resolution frequencies, homogeneous reconstructions carefully average individual frequencies across a large number of images, thereby averaging out the noise. Ideally, one would like to do the same in heterogeneous reconstruction, but that averaging comes with a trade-off as each image may come from a slightly different conformation. Here lies the delicate trade-off at the heart of every heterogeneous reconstruction algorithm: aggregating images is necessary to overcome the noise but degrades the amount of heterogeneity captured. When the mapping from latent space to volume space is parametric, this choice is made implicitly by the parametrization, often by imposing some degree of smoothness. We adopt a scheme that adaptively estimates the optimal number of images to average to reconstruct different frequencies (in the Fourier domain) of different parts of the volumes (in the spatial domain) under the framework of kernel regression.

This is done in four steps: first, a sequence of 50 different kernel regression estimates of the same volume is computed by varying the number of images considered. Second, each estimate is decomposed into smaller subvolumes, and each subvolume is further decomposed into their frequency shells. Third, the best frequency shell of each subvolume is selected automatically by cross-validation and finally, all sub-estimates are recombined to form a full volume. See fig. 1(c) for illustration and appendix A.6 for details.

Surprisingly, despite the relative simplicity of this scheme when compared to black-box deep-learning methods, both our benchmarks in section 1.6 and independent extensive benchmarks [21] find that it outperforms deep-learning methods across datasets with varying SNR, types and complexity of heterogeneity. An additional benefit of this scheme is its transparency: all features of the volume produced can be easily traced back to the images that produced them.

The reprojection scheme and the kernel regression scheme highlight two different interpretations of linear subspace methods. While the reprojection scheme might be interpreted as explicitly modeling heterogeneous distributions as linear combinations of volumes, the kernel regression scheme underscores an alternative and sometimes more appealing interpretation: it merely estimates the orthogonal projection of conformational states onto a low-dimensional subspace. PCA identifies the optimal subspace and endows the resulting embedding with desirable properties: distances and uncertainty are preserved between volume and latent space up to truncation error in the subspace. We next show how to leverage these properties to estimate the conformational density.

### 1.4 Estimating the density of states

Through Boltzmann statistics, the density of a particular state is a measure of the free energy of that state; thus it is a great promise of cryo-EM heterogeneity to accurately recover this density. ^5^ However, estimating volume densities from embeddings obtained by cryo-EM heterogeneity pipelines presents two challenges. First, embeddings distort the space; thus the density of the embedding can be entirely unrelated to the density of the underlying conformational distribution, see [26]. Second, the predicted labels 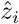 are very noisy, an unavoidable consequence of the low SNR of cryo-EM images; thus the distribution in the embedding reflects not only the distribution of the states but also the noise. As a result, the observed density cannot generally be used to infer the density of the conformational density.

In this regard, the PCA embedding offers favorable properties compared to nonlinear ones: it finds a minimum distortion embedding [1], and the effect of the noise on the embedding is tractable to model. We exploit these properties to recover the conformational density from the PCA embedding. We distinguish two densities: the underlying *conformational density* as the density of unobserved volumes *D*(*x*) (and its representation in PCA space *d*(*z*)), and the *embedding density*, which is the density of the observed predicted labels 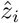 in latent space. Specifically, we consider the kernel density estimator [51] of the predicted labels 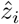:

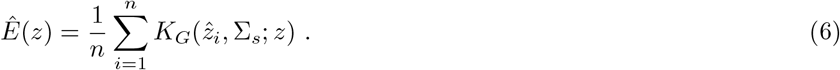

where *K*_*G*_(*µ*, Σ; *z*) is the probability density function of the multivariate Gaussian with mean *µ* and covariance Σ evaluated at *z*, and Σ_*s*_ is set using the Silverman rule [51]. It can be shown that, under some assumptions stated in appendix A.7, the embedding density and the conformational density are related as follows:

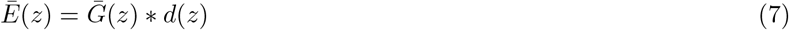

where 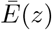 is the expectation of the embedding density 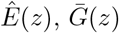 is the expectation of 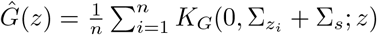 which we name the *embedding uncertainty*, and ∗ denotes convolution. This embedding uncertainty depends on the covariances 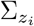 of the latent space labels 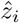, and the kernel width Σ_*s*_. Using this insight, we form an estimator for the conformational density by first estimating the embedding density *Ê* (*z*) and the embedding uncertainty *Ĝ* (*z*), and then use them to estimate the conformational density, see fig. 1(b) for illustration and appendix A.7 for details. Next, we describe how to leverage this conformational density to compute low free-energy trajectories.

### 1.5 Motion recovery

Traditionally, linear subspace methods attempt to recover motions by moving along straight lines in the latent space, corresponding to linear interpolation in volume space. For instance, in 3DVA [38], individual principal components are traversed to generate axes of motion. However, this straightforward linear approach falls short because even a simple linear motion of atoms translates to a highly nonlinear trajectory when observed in volume space. Consequently, a linear path in volume or latent space cannot adequately capture rigid motion, let alone general atomic movements. Fortunately, linear functions locally approximate smooth functions well, allowing small motions to occasionally be captured by traversing a single principal component. In most cases, multiple principal components are needed to accurately represent a single motion, which can lead to significant artifacts when using only one, as illustrated in fig. A.12. Furthermore, multiple axes of motion are typically embedded in the same principal component, as we illustrate below. This lack of separation is clear from the theory: independent variables are not encoded in different principal components by PCA. Alternative decompositions such as Independent Component Analysis (ICA) [8] have that property but require estimating higher-order statistics.

We adopt an alternate strategy based on physical considerations: molecules display stochastic motions, randomly walking from one state to neighboring ones with probability depending on the rate of change of free energy, generally preferring to move towards low free energy states. A physically more meaningful way to estimate trajectory would thus be to find the most likely trajectory between two states. However, extracting accurate free energy profiles can be challenging (see [56]) thus we instead propose a simpler heuristic that overcomes the clear pitfalls of linear trajectories: we generate non-linear motions by computing short and high-density continuous trajectories of volumes between two predetermined conformational states. Since high-density states correspond to low energy, we expect this trajectory to behave like the most likely one. Furthermore, exploiting density allows the trajectory to stay in meaningful regions of latent space.

Given that distances and density are approximately preserved between volume and latent space due to the PCA embedding, we can directly approximate these trajectories in the latent space. Specifically, we compute a trajectory between two states that minimizes the cumulative inverse density. That is, we find the trajectory *Z*(*t*) : ℝ^+^ → ℝ^*d*^ that minimizes the expression

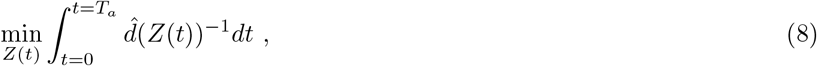

subject to 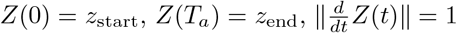 with *T*_*a*_ = min {*t* | *Z*(*t*) = *z*_end_} and 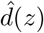 is the estimated conformational density defined in appendix A.7. Minimizing eq. (8) falls under the category of classical optimal control problems, see e.g. [3, 49], and appendix A.8 for our implementation details. Next, we demonstrate on real and synthetic datasets that this approach can recover large and intricate motions.

### 1.6 Validation on a realistic synthetic dataset

We benchmark RECOVAR on a complex synthetic dataset with realistic factors such as SNR, noise distribution, image contrast variation, and motion patterns, see fig. 2(a). This dataset comprises 300,000 images and is designed to mimic the heterogeneity observed in the precatalytic spliceosome complex as reported in previous studies [37, 32]. The simulated complex comprises three bodies that move independently, creating multidimensional heterogeneity. We construct two types of motions: one two-dimensional landscape of top-down motions where two bodies move independently (80% of images) and one trajectory of left-right motions of both bodies (20% of images). Among top-down motions, one intricate trajectory is simulated to have a much higher density. Our goal is to compare the resolution of volumes produced by RECOVAR against established methods and to evaluate its ability to recover density and low free-energy trajectory in realistic conditions.

**Figure 2.**
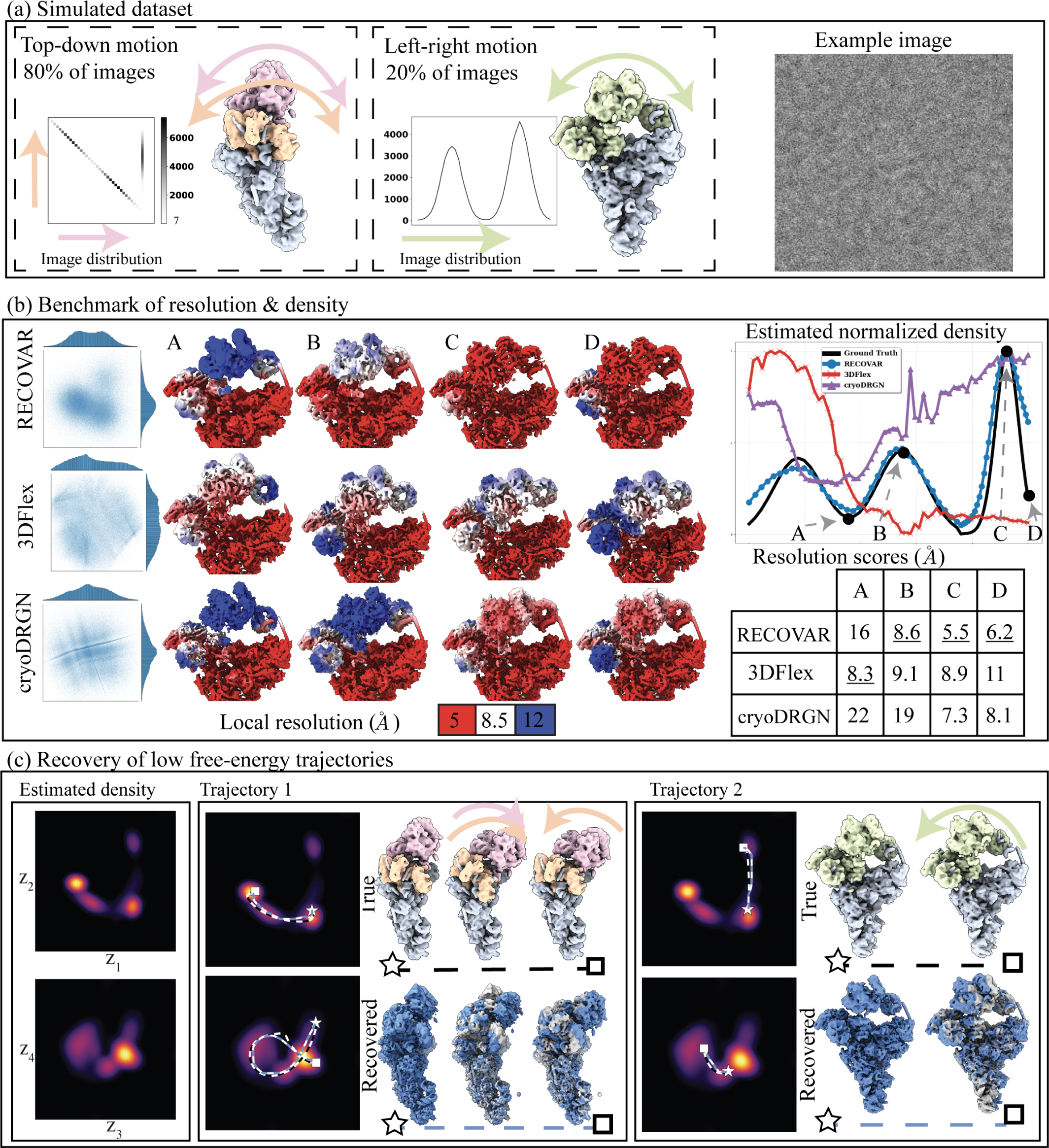
Validation of RECOVAR on a synthetic dataset. **(a)** Simulated dataset of the precatalytic spliceosome with three bodies moving independently. A landscape of motions consisting of 80% images where the orange and pink bodies move independently in top-down motions, including a high-density trajectory where both bodies first move down together, and the orange body then locks back into place. A second motion, consisting of 20% of images, shows the two bodies’ left-right motion. **(b)** Benchmarking of RECOVAR against two deep-learning methods 3DFlex and cryoDRGN. The embedding along two PCs of the embedding is shown for each method, and 4 reconstructed states are plotted in their local resolution. The 90% percentile of the local resolution for each state is shown in the bottom right. The estimated density along the high-density top-down motion of each method is shown in the top right. **(c)** (Left) The estimated 4-dimensional density of the RECOVAR embedding is displayed over pairs of dimensions by integrating the remaining dimensions. (Middle and right) The two ground truth (in black) and two recovered (in blue) trajectories are shown in volume and latent space. The preceding state along the trajectory is overlaid in gray for comparison. The star and the square represent the endpoints.

**Figure 3.**
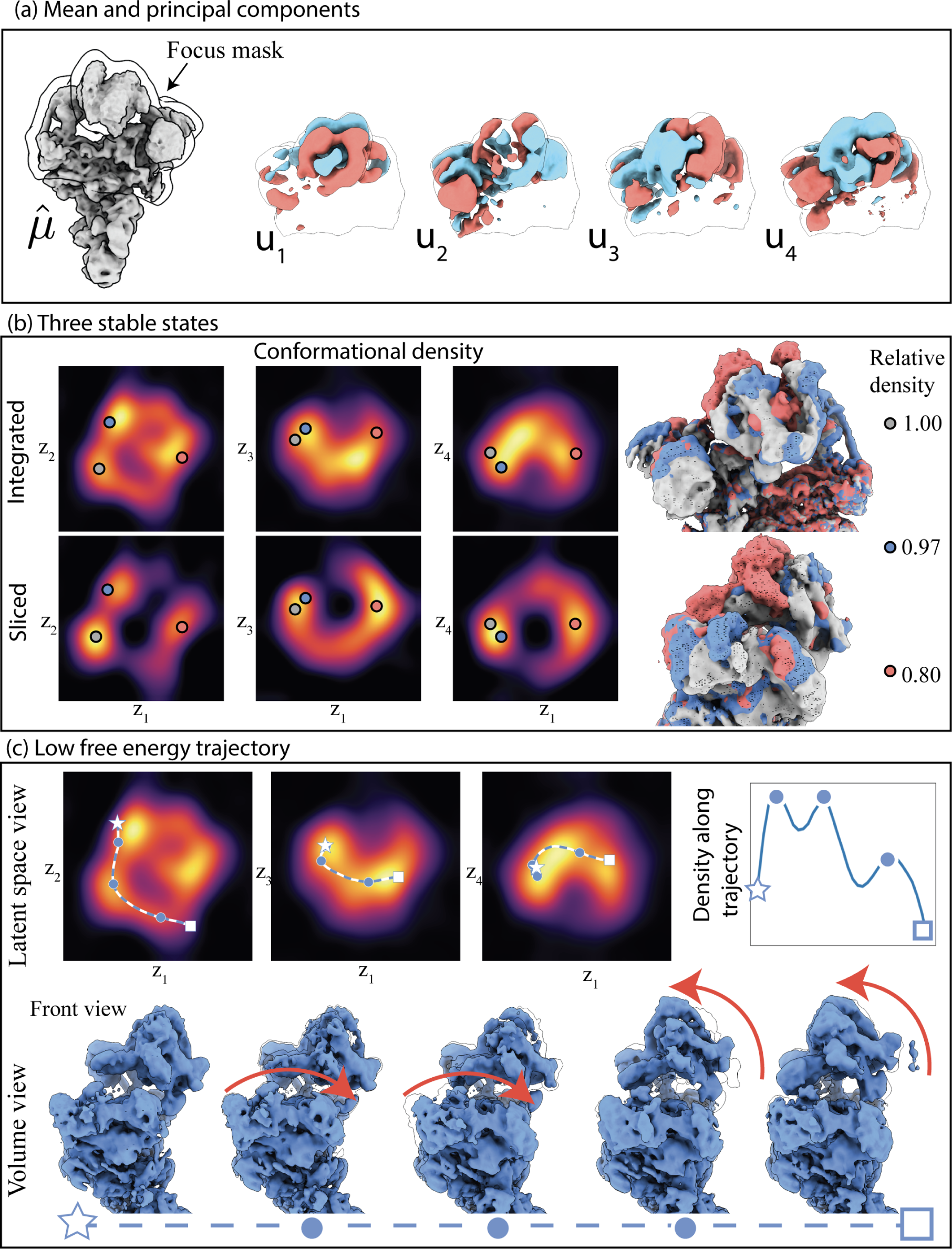
Result of the RECOVAR pipeline applied to the precatalytic spliceosome dataset (EMPIAR-10180). **(a)** Computed mean, with focusing mask overlaid and first four principal components computed. **(b)** Visualization of the estimated conformational density, displayed along pairs of axes displayed by integrating the remaining dimensions (Integrated), and fixing the remaining dimensions to the middle of the grid (Sliced). The three stable states, identified from the local maxima of the conformational density are highlighted. **(c)** An estimated low free energy trajectory shown in latent space and volume space. The trajectory shows two steps: first the arm region moves down then the head region moves up. The trajectory passes near the three local maxima.

We run RECOVAR with a focus mask on the head and arm region and a 4-dimensional embedding. We also benchmark the voxel-based deep-learning method cryoDRGN [65] and the deformation-based deep-learning method 3DFlex [39]. For both deep-learning methods, we perform a hyperparameter sweep^6^ and report the best results. We measure the resolution of states using the 90th percentile of the local resolution. The local resolution is computed by local FSCs between the ground truth and estimated states in a tight mask. Since a large part of the protein is static, as is typical in real datasets, the 90th percentile captures the resolution of the moving part of the molecule.

We evaluate the resolution at four states along the low free-energy trajectory of up-down motions, displayed in fig. 2. We find that in medium to high-density states (B,C,D), RECOVAR outperforms cryoDRGN and 3Dflex at reconstructing high-resolution volumes. In particular, at the highest resolution state (state C), the resolution of RECOVAR is 5.5*Å* compared to 7.3*Å* for cryoDRGN and 8.9*Å* for 3DFlex. On the other hand, in the state with very few images (state A), 3DFlex outperforms RECOVAR and cryoDRGN, which may be due to the ability of 3DFlex’s deformation prior to correctly interpolate in a region where few images are present. These results are consistent with independent extensive benchmarks [21] showing that RECOVAR outperforms 3DFlex, cryoDRGN, and other deep and non-deep learning methods at recovering high-resolution volumes in nearly all cases. Qualitatively, RECOVAR and cryoDRGN volumes have similar properties: low-resolution states display visual features recognizable as low resolution (either no high-frequency features or high-frequency features consistent with noise). In contrast, 3DFlex displays features visually consistent with high-resolution volumes even when the estimate is low resolution (e.g., state D). This is a noteworthy feature of deformation models: the fact that a reconstructed motion visually looks high-resolution does not mean that the result is an accurate reconstruction of the underlying motion, or even that the motion exists at all. This also explains why the halfmap FSC scores of 3Dflex vastly overestimate the true resolution, see fig. A.10 for illustration, and [47] for further examples of bias in deformation models.

We also show kernel density estimates of the cryoDRGN and 3DFlex embeddings, compared to the ground truth and the estimated conformational density of RECOVAR^7^. As expected from theory, the density estimated from cryoDRGN and 3DFlex bear no relation to the true underlying density, but RECOVAR qualitatively reconstructs the true density. The small discrepancy between the true density and the one predicted by RECOVAR is attributed to the inexact forward model used in the density estimation which assumes that the heterogeneity is exactly low-rank.

Despite the highly structured simulated distribution, the raw embeddings for all three methods show no clear trajectories or clusters due to the noisy latent labels, as often observed in real datasets with only conformational heterogeneity. Despite this lack of clear structure in the embedding, the conformational densities estimated by RECOVAR show clear stable states (regions of high densities) and trajectories between them (see fig. 2(c)). This highlights the strength of the statistical framework presented here: even when the raw embedding appears featureless due to the high level of noise in individual images, the careful modeling of the noise on the whole distribution of images allows us to still recover the underlying density, which greatly improves the interpretability of the latent space. Using this estimated density, RECOVAR accurately recovers the intricate low free-energy motion by identifying high-density trajectory as shown in fig. 2(c).

This example also makes it clear why individual principal components as done in [38] should not be interpreted as axes of motions: each motion spans all principal components, and two motions in orthogonal directions are partially embedded into the same principal component directions.

In fig. A.11, we also show the result of all RECOVAR, 3DFlex, and cryoDRGN on a synthetic homogeneous dataset to examine the robustness of these methods to hallucination.

### 1.7 Large motion of the snRNP Spliceosome complex

We apply RECOVAR to the snRNP spliceosome complex (EMPIAR-10073 [33]). We use non-uniform refinement in cryoSPARC [41] to obtain consensus reconstruction, and filter the dataset by running RECOVAR with 20 dimensions and a loose mask to reject compositional heterogeneity, resulting in a stack of 91,763 images. Finally, we apply RECOVAR with a focus mask on the arm, using 4 PCs. Each step of the pipeline is illustrated in fig. 1. The estimated conformational density in fig. 1(d) displays a clear trajectory in the shape of a “U” in the first two PC dimensions. This is in stark contrast to the cryoDRGN embedding shown in fig. A.13 which appears featureless.

We observe one clear maximum in the estimated conformational density, corresponding to a single stable state of the complex, and the density decays monotonically on either side of the stable state. We pick two points at either side of the latent space and produce a low-free energy trajectory transition, showing a large and consistent up-down motion of the arm (fig. 1(d)) that does not have the artifacts typically seen in PCA methods. Other, smaller, left-right motions are best observed in the PCs 3 and 4.

### 1.8 Stable states of the precatalytic splicoseaume

We apply the RECOVAR pipeline to the the precatalytic spliceosome dataset [37] (EMPIAR-10180), which is a well-characterized dataset commonly employed for benchmarking continuous cryo-EM heterogeneity methods. We use the particle stack of 139,722 selected images in [65] downsampled to a size of 256×256, and run RECOVAR with 4 dimensions and a focus mask on the head region of the complex.

In this case, we observe three clear local maxima in the estimated conformational density function, which indicates that while this complex has a large range of seemingly unstable motion, it has in fact multiple stable states.

We pick two endpoints with different positions of the arm and head regions of the complex and generate a low free-energy trajectory from latent space. Interestingly, the low free energy trajectory passes through the three different stable states. The motion appears to comprise two stages: in the first the arm lowers, while the head is static, and in the second stage the head region displays an upward motion. This indicates that while there are states where both the arm and head move concurrently, these are not the most likely states in the dataset and thus not the lowest free-energy transition.

### 1.9 Compositional and conformational changes in an integrin dataset

We apply RECOVAR with 20 dimensions and a loose mask to the *αV β*8 integrin dataset [6] of 84,266 particles. Despite the particle stack being obtained by heterogeneous refinement, we observe a lot of residual compositional heterogeneity, clearly observable from the clusters in the U-MAP visualization of the 20-dimensional embedding. After filtering the dataset of this compositional heterogeneity, we run again RECOVAR with 4 dimensions with a focus mask on the highly flexible arm of the complex. This time, both the PCA and UMAP visualization of the focused analysis reveal no clear structure to latent space.^8^

The estimated conformational density, however, shows two local maxima, with the smaller local maximum corresponding to further compositional heterogeneity that does not seem to be reported in the original study [6]. These results highlight that, even when the SNR is too low to correctly classify individual particles into different clusters, the statistical model of RECOVAR can still detect the presence of a bimodal distribution due to compositional heterogeneity; thereby allowing for better interpretation of the latent space. Aside from the compositional heterogeneity, we also recover the motions of the arm, which shows a wide range of motions.

### 1.10 High resolution states of the ribosome complex

We next analyze a ribosomal subunit dataset (EMPIAR-10076) [7], known for its significant compositional heterogeneity. This dataset has been extensively studied, revealing 14 distinct assembly states through repeated 3D classification [7] in the original research and one additional state identified by cryoDRGN [65].

We run RECOVAR with 20 dimensions on the full particle stack of 131,899 images, and compare it to cryoDRGN (50 EPOCHS) in fig. 5. We display the UMAP of both embeddings, with colors indicating the original classification. The agreement between the clusters and the colors for both methods highlight that RECOVAR and cryoDRGN are both capable of representing this rich heterogeneous distribution of states; underscoring that high-dimensional linear subspaces have large representation power.

While the two methods have similar embeddings, the volumes they recover are different. RECOVAR more accurately estimates the 30S subunit of the 70S ribosome (state A) and recovers state B to higher resolution, evidenced by a more apparent chain structure. This result is consistent with benchmarks showing that RECOVAR outperforms deep-learning methods at recovering high-resolution volumes [21]. We also apply the RECOVAR volume generation scheme to the cryo-DRGN embedding (denoted “hybrid” in fig. 5) and observe that this hybrid scheme outperforms the decoder network in both cases. Therefore, RECOVAR may also be used to improve the resolution of other heterogeneity methods.

Finally, we run RECOVAR on the small cluster of images identified as the 70S (1563 particles). Even in this very small number of images, RECOVAR can identify two stable states of the 70S (blue and purple), corresponding to a rotation of the small subunit relative to the large subunit, with the purple state appearing more stable than the blue state.

## 2 Discussion

### 2.1 Related work

We introduced a heterogeneity analysis pipeline based on PCA with a novel automatic regularization technique and improved efficiency, enabling the fast and robust computation of large numbers of principal components at high resolution. We described an adaptive kernel regression scheme to reconstruct volumes from embeddings. Finally, we leveraged the favorable properties of the PCA embedding to estimate the underlying conformational density from the embedding density by modeling the effect of the noise on the PCA embedding. This ability to accurately estimate the conformational density greatly improves the interpretability of the latent space by enabling the identification of stable states and low free energy motions from their high density.

Several linear subspace methods have been proposed for cryo-EM heterogeneity, including bootstrapping from 3D reconstructions [34, 36, 35, 64, 63, 17, 16], covariance estimation methods [23, 2], probabilistic PCA (PPCA) methods [55], optimization-based methods [38], and methods employing molecular dynamics [59]. We detail the different properties of some of these methods in appendix A.9. However, linear subspace methods have often been thought to be low-resolution, or only able to capture simple distribution or small motion. In contrast, our PCA method coupled with a novel kernel regression scheme produces higher resolution volumes than all other tested methods in extensive benchmarks across SNR, type, and complexity of heterogeneity [21]. Furthermore, we showed that linear subspace methods can capture very rich distributions by computing large dimensional subspaces, and large motions by computing nonlinear trajectories in PCA space.

Another popular class of method for heterogeneity analysis is deep-learning methods, which have received a lot of research interest following the success of cryoDRGN [65] at reconstructing a wide range of heterogeneous distributions (see [11] for a recent review). They can, in principle, represent the distribution with fewer degrees of freedom natively, though it may be preferable not to do so since the overparameterization may be beneficial to train the network [65]. Intuitively, it may seem that this more compact representation may regularize the latent space and thus would lead to higher resolution states. However, the less compact PCA embedding presented here results in higher resolution volumes than deep-learning methods, which may indicate that current variational auto-encoder architectures are under-regularized in other ways, and struggle to distinguish between noise and heterogeneity at low SNR.

The drawback of deep-learning methods for heterogeneity analysis is their lack of mathematical foundation and black-box nature, making interpretation, validation, and hyperparameter tuning particularly difficult. Furthermore, the lack of explicit regularization and metrics for heterogeneity can make identifying suitable parameters difficult, and the user must resort to repeated, expensive computations and subjective visual inspection. This is particularly problematic for methods that explicitly model the deformation as they can produce hallucinated but realistic-looking motions from pure noise or model mismatch (e.g., the presence of residual compositional heterogeneity as in fig. 4).

**Figure 4.**
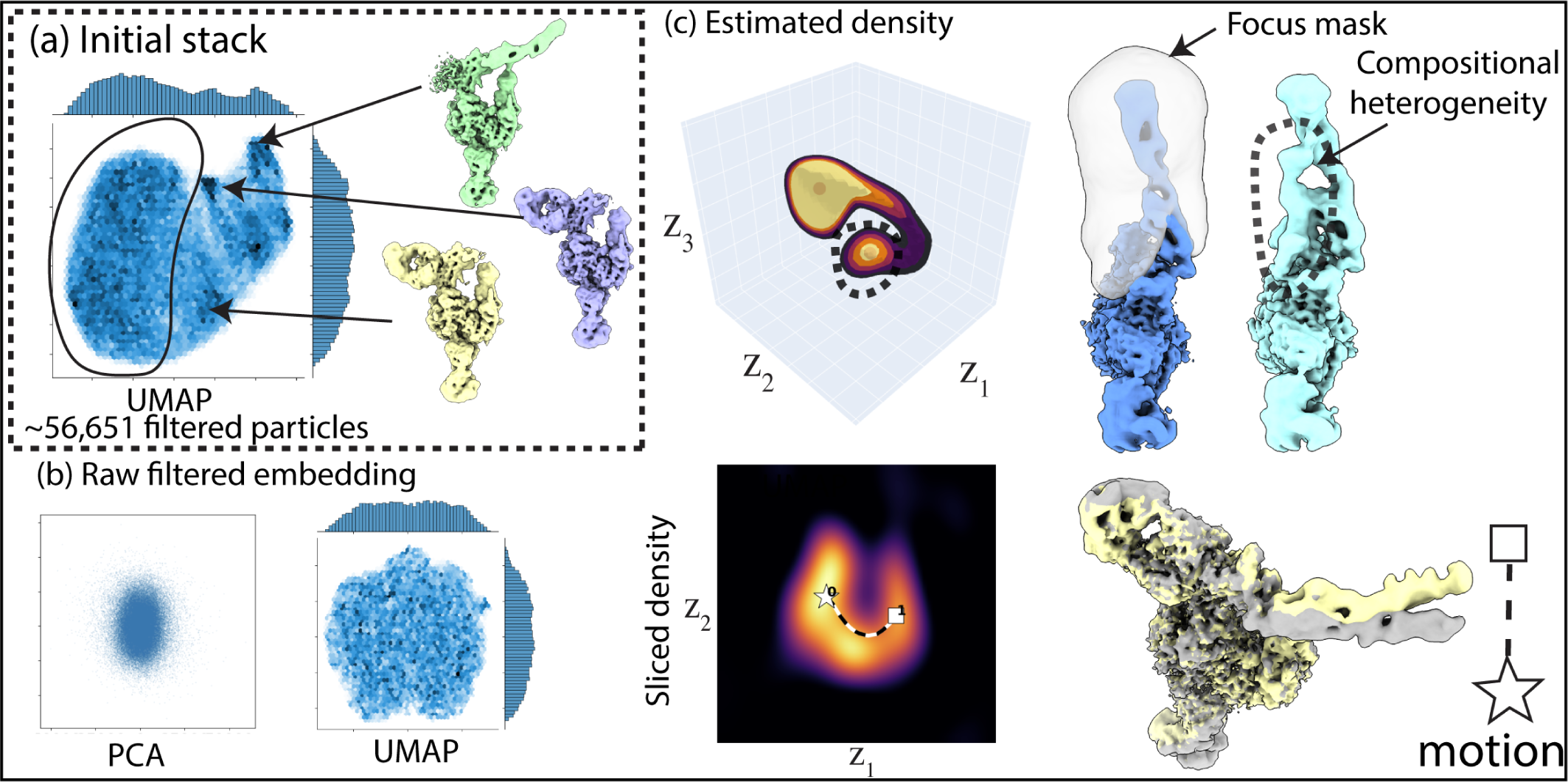
RECOVAR applied to an integrin dataset (EMPIAR-10345). **(a)** Initial run of RECOVAR, showing clusters in the UMAP visualization that correspond to large compositional heterogeneity. **(b)** Visualization of the embedding of the filtered dataset, showing no clear structure (similar to the cryoDRGN embedding in fig. A.13). **(c)** Estimated conformational density, showing a bimodal distribution. The smaller mode corresponds to further compositional heterogeneity. Also pictured: a motion of the arm, plotted in latent space over the density sliced at the mean of the end and start point, and the reconstructed volumes.

In contrast with deep-learning methods, RECOVAR is based on classical statistical and applied mathematics methods. As a result, it is more interpretable, and it inherits crucial properties that are lacking in deep-learning methods: they do not preserve distances or distributions, uncertainty estimates are unavailable, and deep-learning embeddings exhibit unexpected behavior [12]. Consequently, the interpretation of deep-learned latent spaces is challenging, and there is currently no clear way to leverage these embeddings to estimate the conformational density as done in RECOVAR.

### 2.2 Validation of heterogeneity analysis methods

Emerging methods for heterogeneity analysis in cryo-EM encompass a wide range of techniques, from deep learning, manifold learning, and PCA to deformation fields and molecular dynamics. Each approach has its strengths, but a notable challenge arises when validating and comparing the outcomes of these diverse techniques. While all heterogeneity methods yield probability distributions of conformational states as output, the challenge stems from their differing representations and coordinate systems. These differences make direct comparisons a complex task.

Even comparing individual reconstructions is not straightforward when generated by heterogeneous algorithms. As highlighted in appendix A.10, the FSC score, ubiquitously employed to evaluate the resolution of homogeneous reconstructions, can be misleading when applied to volumes obtained by heterogeneity analysis. One reason is that different conformations within a heterogeneous dataset are often highly correlated. Thus, using a correlation-based score to evaluate the reconstruction of a specific state can inadvertently reflect the homogeneous component of the reconstruction. Furthermore, as depicted in the fig. A.10, the “gold-standard” FSC computed from halfmaps is a biased estimate of the FSC between the estimate and the true volume when applied to heterogeneous reconstructions. As a result, the gold-standard FSC and its local FSC analogues, do not accurately reflect the quality of the estimated map.

Covariance estimation and linear subspace methods offer promising avenues to address the challenge of quantifying the consistency of heterogeneous reconstructions across different techniques. First, sampling from each method’s distributions makes it possible to estimate the covariance of each method’s output and compare it with the covariance computed directly from minimally processed images as in appendix A.1. Furthermore, the covariance estimates can be compared using the covariance FSC proposed in appendix A.3. Second, the mapping from volume space to latent space *x* →*U* ^*∗*^(*x*−*µ*) provided by linear subspace methods can embed volumes obtained by different algorithms into the same coordinate system. This mapping could allow for comparisons and alignment of different distributions of volumes obtained through other heterogeneity methods.

### 2.3 Limitations and opportunities

The algorithms in RECOVAR are entirely transparent and relatively simple: the embedding is done by PCA (among the simplest embedding methods), volumes are generated by averaging images, and motions are computed by following highdensity regions of latent space. Each of these steps has opportunities for further improvement: more sophisticated prior in the PCA computation (e.g., using sparsity priors [67]), polynomial kernel regression [18] to generate volumes, and further improvement in the computation of density (e.g., [60]). The fact that RECOVAR already outperforms other methods in several aspects without these improvements provides exciting opportunities for the future development of heterogeneity methods.

A shortcoming of the method presented here is the lack of an automatic choice of the optimal number of principal components for downstream tasks. Since our method demonstrates resilience against overestimation, we recommend using 20 dimensions for exploratory analysis and dataset cleaning, and 4 dimensions for density estimation. More granular analysis can be done using the decay of eigenvalues (e.g., the Scree test [22]) and visualizing principal components and their embeddings. Fortunately, our method computes embeddings of different sizes at virtually no extra time, and thus, we calculate the embeddings with *d* = 1, 2, 4, 10, 20 in our implementation by default. We note that related theory predicts the optimal choice based on observation count and noise levels [9] but does not cover prediction for the method presented here. This area presents an opportunity for theoretical exploration.

A limitation of our current density estimation method is its inability to estimate density over a large number of principal components; this is a purely computational problem: the run time of our deconvolution method in appendix A.7 is exponential in the number of principal components. We emphasize that this is a shortcoming of density estimation, not of linear subspace methods, and we expect that recent methods for high dimensional density estimation [60] might help resolve this issue. In the meantime, we advocate for the use of a focusing mask to focus the heterogeneity on a zone of interest while factoring out other sources of heterogeneity to reduce the required PC dimension. Further improvement in density estimation could also be achieved by modeling the uncertainty in pose estimation as in [56].

Thanks to the white-box nature of the pipeline, the different components of the RECOVAR can be easily integrated into different frameworks which presents many opportunities to pick and choose the best components between different heterogeneity methods. For example, we showed in fig. 5 that RECOVAR’s kernel regression scheme can be applied to the cryoDRGN embedding and improve the resolution of cryoDRGN’s volumes. Similarly, the density estimation and motion trajectory could be used in conjunction with methods based on molecular dynamics such as [59].

**Figure 5.**
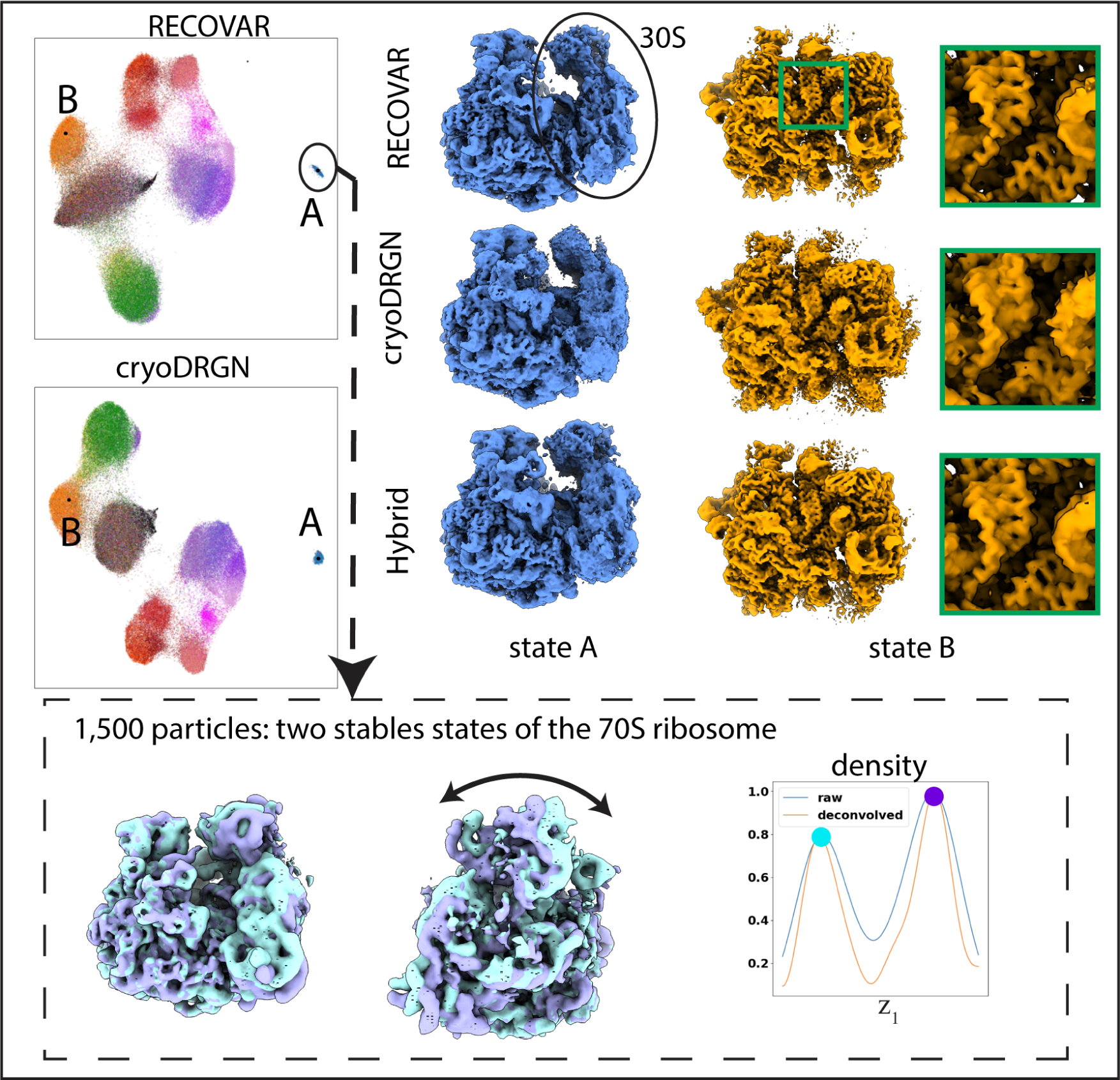
Result of RECOVAR on ribosomal subunit dataset (EMPIAR-10076). **(Top)** Embedding of RECOVAR and cryoDRGN, and two states produced by each method. All volumes are sharpened with B-factor −85*Å*^2^, except state A of cryoDRGN for which the 30-S subunit is less visible when sharpened. RECOVAR better resolves the small subunit and recovers a higher resolution reconstruction of the B-state. **(Bottom)** RECOVAR run on 1,500 particles identified as the 70S ribosome. Two stable states are clearly identified, corresponding to a rotation of the small subunit.

The introduction of the Bayesian framework was central to the resolution revolution in cryo-EM [24] and has dramatically contributed to its democratization thanks to its robust parameter estimation framework that requires relatively little user input. In contrast, the current state of heterogeneity analysis typically involves ad-hoc filtering, hand-tuning of various parameters, and the prospect of drawing false conclusions from hallucinated motions. The framework presented here generalizes the Bayesian framework used for homogeneous reconstruction and thus offers similar potential in robustly and confidently reconstructing wide ranges of heterogeneous distributions from cryo-EM datasets with little need for tuning.

The Bayesian framework was particularly critical in developing robust homogeneous methods that infer poses and volumes. These algorithms typically alternate between predicting poses and predicting volumes. However, poor assignment of poses produces inaccurate volumes, and inaccurate volumes, particularly from overfitting, also cause poor alignment. The significant advance of the FSC regularization introduced in [45] was to resolve this issue: thanks to the adaptive regularization recomputed at each step, resolution and alignment can progress slowly together. Our regularization strategy generalizes this scheme, and our method’s high efficiency allows it to be run repeatedly within an alternating framework. Therefore, it is a promising method to address the problem of jointly inferring conformations and poses. Hence, we expect that future work using this framework will result in more accurate pose estimation for highly heterogeneous datasets, thereby improving the input to all heterogeneity reconstruction algorithms.

## A Methods

### A.1 Estimation of the mean and covariance of the conformational distribution

We describe the kernel regression discretization used to solve the least-squares in eq. (2). We denote functions with bold fonts and their discretization on a Cartesian grid without bold font, e.g. a particular conformation, represented in the Fourier domain, is denoted as **x**(*ξ*) : ℝ^3^ → ℂ, and its discretization on a Cartesian grid as 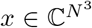 where *x*_*k*_= **x**(*ξ*^*k*^), and {*ξ*^*k*^}_*k*_ is *x* an enumeration of the three dimensional Cartesian grid.

Pixel *j* of the Fourier-transformed image *i* is modeled as *y*_*i,j*_ = *c*_*i,j*_**x**_*i*_(*ξ*_*i,j*_) + *ϵ*_*i,j*_ where *c*_*i,j*_ is the CTF, **x**_*i*_(*ξ*) : ℝ^3^ → ℂ is the Fourier transform of the scattering potential of the conformation present in image *i, ξ*_*i,j*_ ∈ ℝ^3^ is the 3D frequency sampled by pixel *i, j*^9^ (given by the fixed pose *ϕ*_*i*_), and *ϵ*_*i,j*_ is the noise in pixel *i, j*. We consider the following kernel regression estimate for ***µ***(*ξ*), the mean of the distribution of conformations **x**_*i*_(*ξ*):

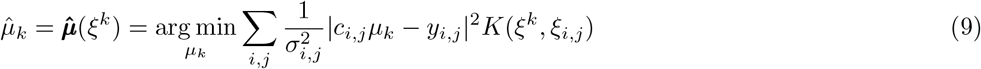

This is the formulation used in RELION, where *K*(*ξ*^*k*^, *ξ*_*i,j*_) is the triangular kernel: 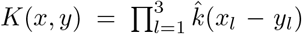 where 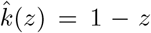 when |*z*| *<* 1 and 0 otherwise. ^10^. This matrix equation can then be efficiently solved in 𝒪 (*N* ^3^ + *nN* ^2^) operations, analogous to the E-step of the E-M algorithm used in iterative refinement as described in [44].

The regularized covariance estimator in eq. (3) can be similarly discretized as:

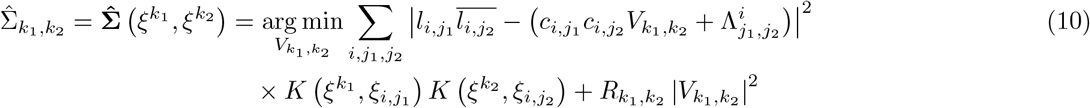

where 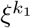 and 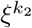 are two grid frequencies, 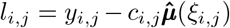 is the deviation from the projected mean in pixel *j* of image 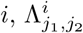 is the covariance of the noise between pixels *j*_1_ and *j*_2_ of image 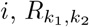 are the regularization weights. We estimate 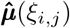 off the grid by spline interpolation. The optimality conditions lead to:

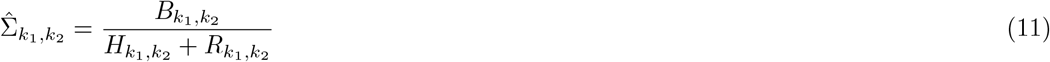

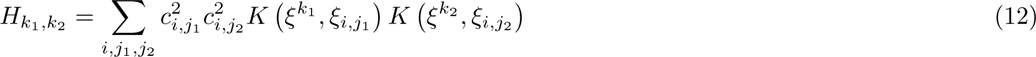

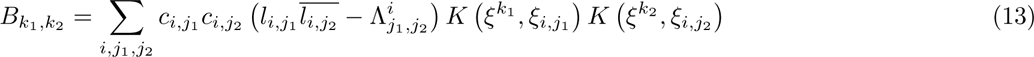

Using this formulation, *d* columns of 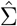 may be computed in 𝒪 (*d*(*N* ^3^ + *nN* ^2^)) in a single pass through the data.

Experimental images often contain multiple particles within each image that can severely bias the estimation of the covariance. To mitigate bias, we use a real space solvent mask to filter out these unwanted particles. To generate per-image masks, we dilate a three-dimensional loose solvent mask and project it using inferred poses for each image. We then threshold and smooth by a cosine-softening to generate mask *M*_*i*_ and apply them to the mean-subtracted images. This masking has the effect of changing the noise statistics of the image, e.g., if the pixels of the original image have noise covariance of Λ_*i*_, then the covariance of the noise of the masked image is *M*_*i*_Λ_*i*_*M*_*i*_^*∗*^, thus we change the noise covariance in eq. (10) accordingly.

This kernel regression scheme allows us to efficiently compute elements of the covariance matrix, but computing all 𝒪 (*N*^6^) elements of the covariance matrix is still prohibitive. This kernel regression scheme allows us to efficiently compute elements of the covariance matrix, but computing all 𝒪 (*N*^6^) elements of the covariance matrix is still prohibitive. We next describe numerical algebra techniques to approximate leading eigenvectors of 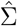 using a subset of entries to address this issue.

### A.2 Approximating principal components from subsets of columns

Having estimated a subset of columns of 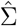, which we denote 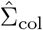, we next estimate the covariance of conformations restricted to the space spanned by these columns. That is, we assume that each volume *x*_*i*_ can be written as 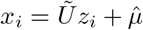 for some *z*_*i*_ where 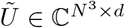 is an orthogonal matrix whose columns spans the columns of 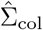 computed from an SVD of 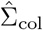^11^, and estimate the covariance of the distribution of *z*_*i*_. A least-squares covariance estimator of *z*_*i*_ (a matrix of size *d* × *d*), similar to the one in eq. (3) but smaller dimensions, may be written as:

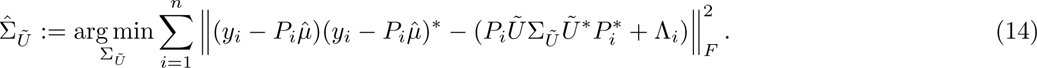

The optimality conditions for this least-squares problem are:

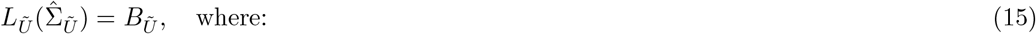

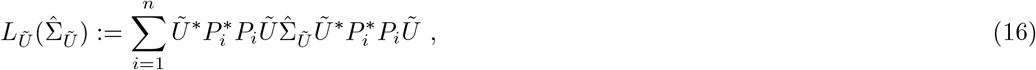

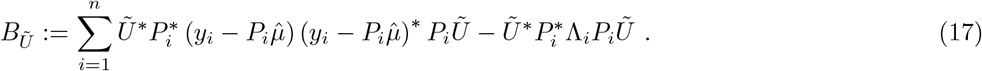

This system of equations can be formed in 𝒪 (*n* (*N*^3^*d* + *d*^4^)) and solved in 𝒪 (*d*^6^) operations.

The strategy of approximating a matrix from a subset of its columns is a fundamental technique in numerical linear algebra and the basis for schemes such as the Nyström extension [61] and the CUR decomposition [29].^12^ The approximation accuracy of these schemes depends on the matrix’s properties and the chosen columns. While near-optimal choices exist in theory (see, e.g. [25, 5]), our case differs due to our goal of computing eigenvectors of the true covariance Σ using noisy estimates with heteroscedastic noise through the estimation problem in eq. (10).

Unfortunately, our case is poorly covered by theory, so we resort to a heuristic scheme that balances two objectives: computing a set of uncorrelated columns, and computing high SNR columns. Nearby frequencies are highly correlated under the bag of atoms model [52, 14], hence we ignore columns within a set distance of columns already picked. We compute an SNR proxy for column *k* as: 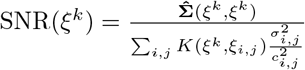 as defined in eqs. (9) and (10). We then make the following greedy choice: first, add all the columns to the considered set. Then, iteratively add the highest SNR column in the considered set to the chosen set, and remove all nearby frequencies from the considered set, until the chosen number of columns is reached. Finally, we take advantage of the Hermitian symmetry property of the covariance matrix 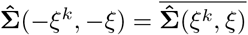 to generate columns corresponding to frequencies −*ξ*^*k*^ for all *ξ*^*k*^ in the selected set. We summarize the proposed PCA method in algorithm 1 and examine its accuracy with different choices of parameters on a simulated dataset in fig. A.6.

#### Algorithm 1

Proposed PCA method

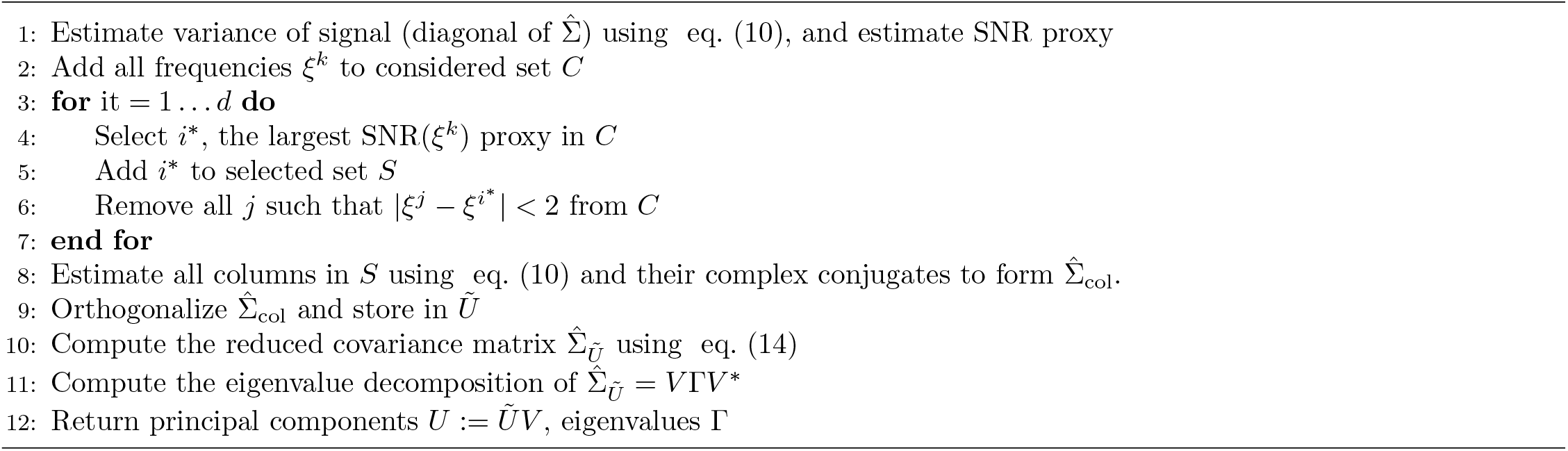

### A.3 A generalized FSC regularization scheme

We introduce a scheme that generalizes the Fourier Shell Correlation (FSC) regularization initially proposed in [44] for homogeneous reconstruction. The foundation of this regularization scheme is rooted in the simplifying assumption that both the signal and the noise are independently and identically distributed within one frequency shell, though they follow different distributions in different shells. This simplification allows us to decouple computations over these shells.

Thus, we consider the signal and noise in a specific frequency shell where the signal *x* follows a Gaussian distribution with zero mean and variance *κ*^2^, while the noise *ϵ* follows a Gaussian distribution with zero mean and variance *σ*^2^. This leads to the simple scalar-valued forward model:

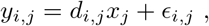

where *x*_*j*_ ∼𝒩 (0, *κ*^2^) represents frequency *j* within the shell, *y*_*i,j*_ is the *i*−th observation of frequency *j* corrupted by noise *ϵ*_*i,j*_ ∼𝒩 (0, *σ*^2^), and *d*_*i,j*_ ∈ ℝ models the CTF. We then estimate the underlying signal *x* by solving the following least-squares problem:

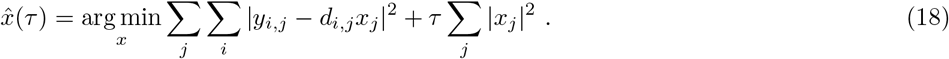

Here, *τ* serves as a regularization weight. The solution to eq. (18) is:

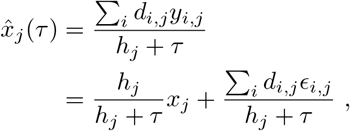

where 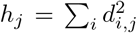. The optimal value of *τ*, which minimizes the mean-squared error (MMSE) is known to be 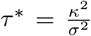. However, estimating *τ*^*∗*^ is our goal. To do this, we begin with an initial value *τ*_0_ and calculate two estimates 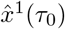 and 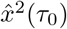 by randomly splitting the dataset into two halves. We then estimate their correlation using the concept from [42]:

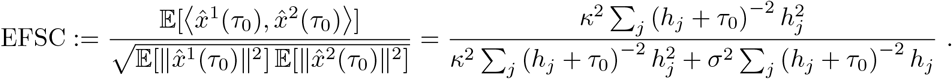

Solving for 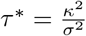, we obtain the estimate:

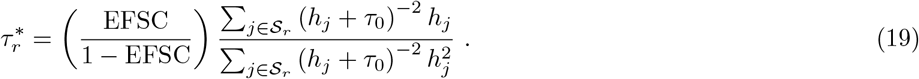

where *r* indexes the specific frequency shell 𝒮_*r*_ = {*j* | *r* ≤ ∥*ξ*^*j*^∥ *<* (*r* + 1). See [20] for a related scheme to compute Wiener weights. For comparison, under our assumptions and notation, the estimate proposed in [45] is equivalent to:

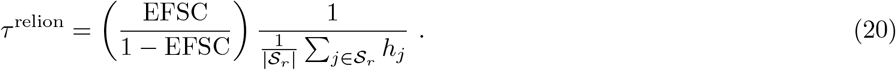

where |𝒮_*r*_| denotes the number of voxels in the *r*-th frequency shell. Notably, if the pose distribution is uniform (more precisely if *h*_*j*_ are constant within a frequency shell), then the two estimates in eq. (19) and eq. (20) agree, but eq. (19) also accounts for the non-uniform case. This new estimator can be applied to homogeneous reconstruction, and we expect it to perform better for datasets with highly non-uniform pose distributions. However, the effect is most pronounced for covariance estimation since sampling of the entries of the covariance matrix in a given column and frequency shell is highly non-uniform even when the pose distribution is uniform.

We use this regularization scheme to set entries of the regularization matrix *R* in eq. (3) by applying it independently in each column. We replace the EFSC with the empirical FSC computed between halfsets, initializing *R* from a multiple of the regularization weights of the mean in eq. (2) and iterating this process 20 times, which has been observed to converge in practice.^13^ That is, we compute the entries of the weight matrix *R* as follows:

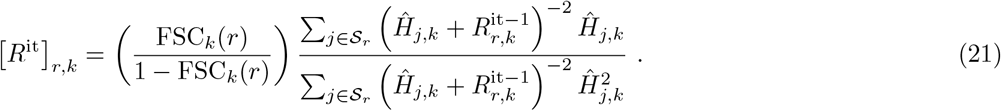

Here, FSC_*k*_(*r*) is the FSC between column *k* of the two covariance matrices 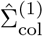 and 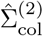 computed from the two half-datasets, and *Ĥ* is the average of the two values of *H* calculated from eq. (12). We perform these computations only for the columns chosen in appendix A.2, indicated by the subscript col. The outlined algorithmic steps are summarized in algorithm 2.

#### Algorithm 2

Computation of the regularization parameters

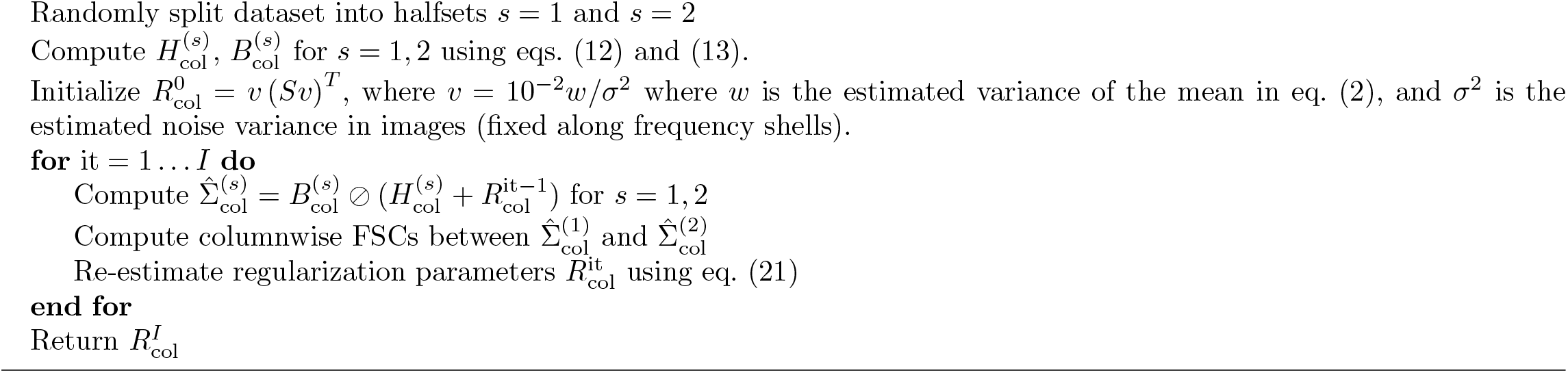

### A.4 Contrast correction in covariance estimation

Cryo-EM images sometimes display variations in contrast. This effect is captured in the image formation model through an additional scaling factor *a*_*i*_ ∈ ℝ^+^:

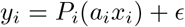

If not accounted for, this contrast variation can manifest as a spurious source of heterogeneity, often displayed in recovered trajectories with appearing or disappearing parts of the volume (see fig. A.7). While some algorithms for consensus reconstruction attempt to infer the scale parameters *a*_*i*_ as part of the algorithm [46], the estimation is poor when significant heterogeneity is present, so they should be estimated as part of a heterogeneous reconstruction. As the contrast is not factored into the covariance estimation in eq. (3), our covariance estimate reflects the covariance of the contrasted distribution of states, which we denote by Σ_*ax*_. Assuming that contrast factors *a*_*i*_ and states *x*_*i*_ are independent, Σ_*ax*_ can be related to the (non-contrasted) covariance of the distribution Σ_*x*_ as follows [50]:

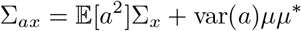

That is, the contrasted covariance differs by an unknown scaling factor close to 1 (e.g., 𝔼 [*a*^2^] ≈ 1.16 in fig. 3) which only scales up the eigenvalues, and is corrupted by a rank-one component var(*a*)*µµ*^*∗*^. We assume that Σ_*x*_ = *U* Γ*U* ^*∗*^ is low-rank and *U* ^*∗*^*µ* = 0^14^, and we recover the original subspace by projecting out the mean component:

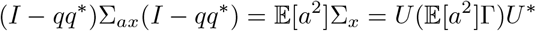

where 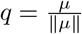. That is, the principal components are correct, but the eigenvalues are overestimated by a small factor which we ignore. See fig. A.7 for an illustration of the effect of the covariance correction step on the estimated principal components and estimated per-image contrast.

Furthermore, the low-rank decomposition of Σ_*x*_ = *U* Γ*U* ^*∗*^ can be efficiently computed from the low-rank decomposition of 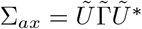 as shown in algorithm 3:

#### Algorithm 3

Update of low-rank decomposition by projecting out mean component

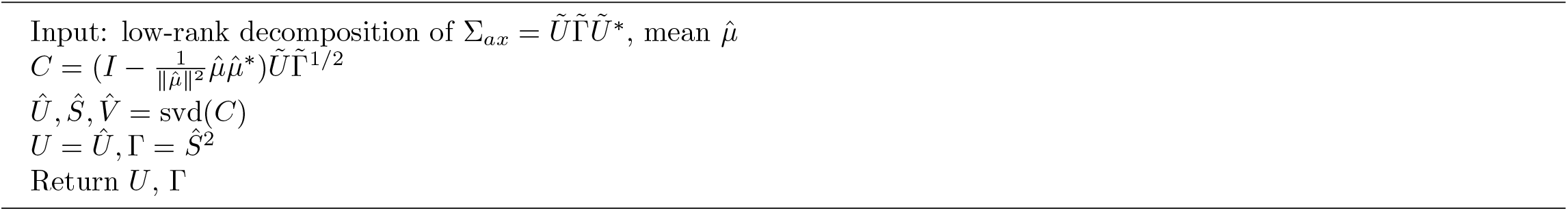

### A.5 Estimation of per-image latent distribution and contrast

We estimate the per-image distribution of likely state and contrast associated with image *y*_*i*_ using a MAP estimate. Assuming *z*_*i*_ follows a Gaussian distribution 𝒩 (0, Γ) and using a flat prior on *â*_*i*_, the MAP estimate is written as the solution of the following optimization problem ^15^:

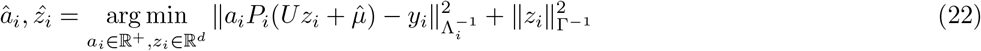

To solve this minimization, we perform a grid search over the contrast variable *a*_*i*_ ∈ [0, 2] and explicitly minimize over the latent variables *z*_*i*_ ∈ ℝ^*d*^. Solving eq. (22) for a fixed contrast *a*_*i*_ involves the normal equations:

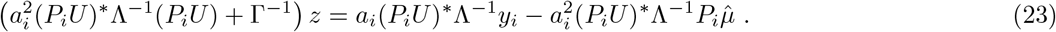

The computational complexity of constructing and solving eq. (23) is 𝒪 (*N* ^2^*d*^2^ + *d*^3^). Therefore, the näive computational complexity of solving this problem for all *n*_*a*_ contrast values of *a*_*i*_ is 𝒪 (*n*_*a*_*N* ^2^*d*^2^ + *n*_*a*_*d*^3^) operations. However, if the matrix (*P*_*i*_*U*)^*∗*^Λ^*−*1^(*P*_*i*_*U*), and vectors (*P*_*i*_*U*)^*∗*^Λ^*−*1^*y*_*i*_, (*P*_*i*_*U*)^*∗*^Λ^*−*1^*P*_*i*_*µ* are precomputed and stored, the complexity drops to 𝒪 (*N* ^2^*d*^2^ +*n*_*a*_*d*^3^). In our experiments, we set *n*_*a*_ = 50, and the additional cost of optimizing over contrast is small compared to the cost of optimizing over *z*_*i*_ alone since the 𝒪 (*N* ^2^*d*^2^) term dominates.

The covariance of the state, denoted as 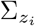, is directly computed from eq. (4) as: 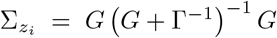 where 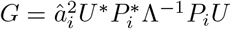 in 𝒪 (*N* ^2^*d*^2^ + *d*^3^) operations per image.

### A.6 Generating volumes from heterogeneous datasets using adaptive kernel regression

We describe in more detail the method presented in section 1.3. Given a fixed latent position *z*^*∗*^, we compute kernel regression estimates of the form:

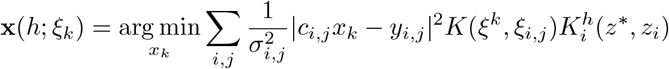

the kernel *K* is the triangular kernel, *h* is the kernel width capturing the number of images used to reconstruct a particular state, and 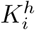 is per-image kernel encoding the confidence of the label *z*_*i*_: 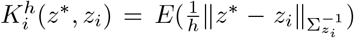 where *E* is a piecewise constant approximation of the Epanechnikov kernel [18]. We choose the Epanechnikov kernel rather than the more common triangular kernel because it is optimal in a mean-square error sense [51] and performs slightly better empirically. Furthermore, we use an approximation of the Epanechnikov kernel rather than the Epanechnikov kernel for computational efficiency, as it allows us to compute many kernel estimates in a single pass through the data. For example, the kernel regression scheme computes a conformation in 8 minutes on one NVIDIA A100 80GB GPU for a dataset of 300, 000 images of size 256^2^.

We compute 50 different estimates by varying the kernel width *h* for a single state *z*^*∗*^ and take the best parts from each by optimizing with cross-validation as follows. Each dataset is split into two sets: from one halfset, we compute the 50 estimates 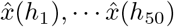, from the other subset we compute a single low-bias, high-variance template 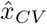 by using a small number of images (100 by default) which are the most likely to be of conformation *z*^*∗*^ according to the statistical model. Each kernel estimate is then subdivided into small subvolumes by real-space masking, and each small subvolume is decomposed into its frequency shells after a Fourier transform. The cross-validation metric we use for each frequency shell is: 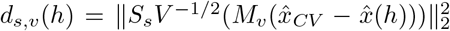, where *S*_*s*_ is a matrix that extracts shell *s, M*_*v*_ is a matrix that extracts the subvolume *v*, and *V* is a diagonal matrix containing the variance of the template. For each frequency shell, the minimizer (over *h*) of the cross-validation score *d*_*s,v*_(*h*) is selected, and a full volume is reconstructed by first recombining all frequency shells of a subvolume, and then recombining the subvolumes. Finally, the volume is B-factor sharpened and regularized by local filtering.

### A.7 Estimating the conformational density

We motivate and elaborate on the scheme to estimate the density in section 1.4. Assuming that the low-rank assumption holds exactly (that is, volumes can be represented as *x*_*i*_ = *Uz*_*i*_ + *µ*), then each image may be represented as:

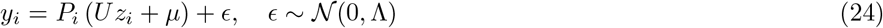

The maximum-likelihood estimate (MLE) of the unobserved *z*_*i*_ given *y*_*i*_ is 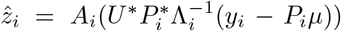 where 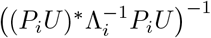^16^. It follows that 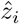 is normally distributed with distribution 𝒩 (*z*, *A*). Assuming that *z* and *A* are independent, this implies that the expected value of the kernel density estimator with kernel width Σ_*s*_ in eq. (6) of the MLE labels is:

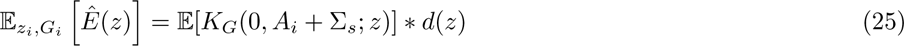

where the *K*_*G*_(*µ*, Σ; *z*) is the multivariate Gaussian probability density function with mean *µ* and covariance Σ evaluated at *z*. We estimate 𝔼 [*G*(0, *A*_*i*_ +Σ_*s*_; *z*)] by Monte-Carlo sampling with *m* = 1000 images chosen at random 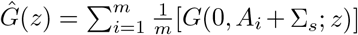, and propose the following regularized estimator for the distribution of *d*(*z*):

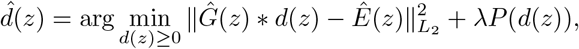

Where 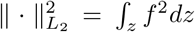, which we discretize on a Cartesian grid, 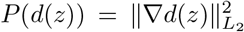 is a prior promoting a smooth distribution and *λ* ∈ ℝ^+^ is a regularization parameter. Picking *λ* correctly is crucial, and we recommend using the L-curve criterion [15] to check that the solution is stable to perturbations in *λ* and to initialization of the optimization algorithm. Our implementation outputs the L-curve and estimates 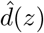 for several values of *λ* so that the user may perform these sanity checks, see fig. A.8 for illustration.

### A.8 Traversing latent space

We generate a motion between two latent space coordinates *z*_st_ and *z*_end_ by solving the minimization problem described in eq. (8). This problem can be approached using dynamic programming by first computing the *value function v*(*z*), defined as:

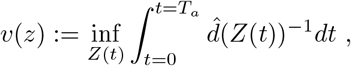

where *Z*(*t*) : ℝ^+^ → ℝ^*d*^ is a continuous trajectory satisfying *Z*(0) = *z, Z*(*T*_*a*_) = *z*_end_ with *T*_*a*_ = min {*t* ≥ 0|*Z*(*t*) = *z*_end_} and 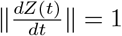. This value function *v*(*z*) is the viscosity solution of the Eikonal equation [3]:

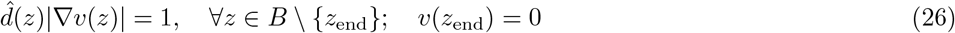

where *B* is the domain of interest. Thus, *v*(*z*) can be computed by solving this partial differential equation. Once eq. (26) is solved, the optimal trajectory can be obtained by finding the path orthogonal to the level curves of *v*(*z*), which can be computed numerically using the steepest descent method on *v*(*z*) starting from *z*_st_.

The Eikonal equation can be discretized and solved using variants of Dijkstra’s algorithm for finding shortest paths on graphs [49]^17^. We choose the domain *B* to be a *d*−dimensional rectangle with lower and upper bounds in dimension *j* equal to the 1st and 99th percentile of the distribution of 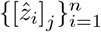. We then discretize the problem by evaluating 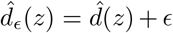 on a *d* dimensional uniform grid. Adding a small constant *ϵ* ensures the existence of a solution and the stability of the numerical method. We set 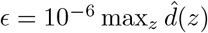.

The computation of the trajectory is dominated by the evaluation of 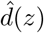 on the grid, resulting in a computational cost of 𝒪 (*g*^*d*^*d*^2^*n*), where *g* is the number of grid points in one dimension which we set to *g* = 50 in all experiments. This limits the applicability of this method to low values of *d*, and therefore, we use this method only in dimensions up to 4. To compute a higher dimensional trajectory, we iteratively increase the dimension of the trajectory by one while keeping the previous dimensions fixed, starting from the 4-dimensional trajectory. This heuristic is not guaranteed to solve the high-dimensional minimization problem, but it performs well in practice.

### A.9 Comparison to other methods for computing linear subspaces

Other popular methods for computing linear subspaces in cryo-EM heterogeneity analysis include PPCA [55] and 3DVA [38]. We briefly summarize them here:

1. PPCA: Find the most likely *U* under the statistical model: *y*_*i*_ = *P*_*i*_(*µ* + *Uz*_*i*_) + *ϵ* where *ϵ* ∼ 𝒩 (0, Λ), and *z*_*i*_ ∼ 𝒩 (0, *I*) for *i* = 1 … *n*. This maximum likelihood problem is typically solved using expectation-maximization (EM), by marginalizing over the variables *z*.
2. 3DVA: Find the minimum of the objective function 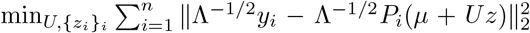. The original paper [38] uses an alternating minimization to find the minimum.^18^
3. Covariance approach: estimate the covariance of the distribution of conformations *x*_*i*_, and find the best low-rank approximation *U* of the covariance matrix.

The covariance approach, which includes RECOVAR, aims to find the low dimensional subspace that best captures the distribution of states given the estimated covariance (i.e. it seeks to minimize the error in the covariance), whereas the other methods compute the subspace that best captures the variation in the dataset of images (i.e. they seek to minimize the residual in image space). These are in general not the same subspace, particularly if there is a highly non-uniform distribution of viewing directions. The difference between 3DVA and PPCA is that whereas PPCA marginalizes over *z* (that is, it uses “soft assignment”), 3DVA optimizes over *z* (that is, it uses “hard assignment”).

3DVA is motivated by the fact that, in the case where *P*_*i*_ = *I* and Λ = *ϵI*, PPCA converges to PCA in the limit of *ϵ* → 0 [43]^19^. However, that statement does not extend to the case where *P*_*i*_ ≠ *I* hence it is not applicable for cryo-EM applications. Therefore, PPCA, PCA, and 3DVA will generate different optimal subspaces.

In the case where the low-rank model is exact, both the covariance approach [23] and the maximum likelihood estimator of PPCA are consistent, thus they will recover the exact subspace as *n*→ ∞. 3DVA enjoys no such theoretical guarantees, and since it effectively treats the noise as signal, 3DVA will compute an optimal subspace to represent both variations in the signal and the variations in the noise. In cryo-EM applications, noise dominates at high resolution, and thus we expect that the subspace recovered by 3DVA will converge to pure noise at sufficiently high resolution. This can explain why 3DVA requires the user to set a low-pass filter resolution, where the SNR is relatively high, and the “streaking” artifacts are sometimes observed in outputs of 3DVA (which may be principal components of the noise). Nevertheless, at sufficiently high SNR and uniform distribution of angles, the three approaches should return similar linear subspaces.

There are three main advantages of the covariance approach compared to PPCA. First, PPCA (and 3DVA) rely on local optimization schemes, which may not converge to the global minimum^20^. Second, the covariance estimator is uniquely defined. This property allows us to estimate the SNR from two independent copies as in appendix A.3. In contrast, the matrices *U* returned by PPCA (and 3DVA) are defined only up to an orthogonal matrix, so a similar approach does not näively apply; though this does not preclude other novel regularization schemes to be crafted for PPCA. The third advantage is the time complexity: the dominant term in the complexity of RECOVAR is 𝒪 (*nN*^2^*d*), whereas the complexity of both 3DVA and PPCA is both 𝒪 (*nN*^2^*d*^2^*s*) where *s* is the number of iterations (20 by default in cryoSPARC)^21^. For large linear subspace dimension *d*, the method presented here is thus much more efficient than 3DVA and PPCA, e.g., for a dataset of 300, 000 images with box size 256 and on the same hardware, RECOVAR computes 100 principal components in 4.5 hours, whereas 3DVA computes 20 principal components in 16 hours, and throws GPU out-of-memory errors at 25 principal components.

PPCA and 3DVA have their advantages: fixing the number of components a priori to a small number may act as an effective regularizer, and the optimization directly over images, rather than the two-step process of the covariance approach, may be preferable for the highest accuracy. E.g., in fig. A.12 3DVA appears to outperform RECOVAR at estimating accurately some of the principal components for some parameter choices at high SNR, likely due to this rank constraint. Furthermore, the flexible iterative framework used in PPCA may offer the potential for future incorporation of more sophisticated priors imposing sparsity or non-uniform resolution.

### A.10 FSC scores for heterogeneity

We illustrate the deceptive nature of FSC scores in determining the resolution of volumes generated by heterogeneous algorithms in fig. A.9. The intriguing observation is that while the reprojected state appears visually as low resolution, the conventional interpretation of the FSC score would lead us to conclude that it is a Nyquist resolution estimate of the true state. On the other hand, despite being visually a much better estimate, the FSC score indicates that the kernel reconstruction (“reweighted”) resolution is around half the reprojected resolution. This puzzling phenomenon can be explained by how the reprojected state is computed: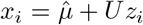. That is, it is a linear combination of the mean and the principal components. The principal components *U* are estimated at a lower resolution than the mean 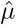 and are regularized adequately, so they do not contribute much noise. Consequently, the state essentially equals the mean in high-frequency shells. Furthermore, the mean is highly correlated to the true state (as the blue curve depicts). Thus, the reprojected state appears highly correlated to the ground truth. While the impact is not as pronounced in the reweighted reconstruction, a similar phenomenon occurs on a more localized scale since any reconstruction based on image averaging necessarily incurs some averaging over different states.

Other methods for heterogeneity reconstruction might not offer the same level of interpretability as the reprojection scheme. However, we anticipate that they would exhibit similar behavior since the static parts of the molecule are more easily resolved to a higher resolution. Adjustments to the FSC calculation, such as the mean-subtracted FSC proposed in [31], or using a local FSC, might provide some relief but could likewise be misleading.

A further, separate issue arises when attempting to estimate the resolution of experimental maps from the “gold-standard” FSC computed from halfmaps. While the gold-standard FSC accurately estimates the true FSC when the dataset and the method are homogeneous, the score can be very biased when applied to heterogeneous reconstructions from heterogeneous datasets, see fig. A.10 for illustration.

**Figure A.6:**
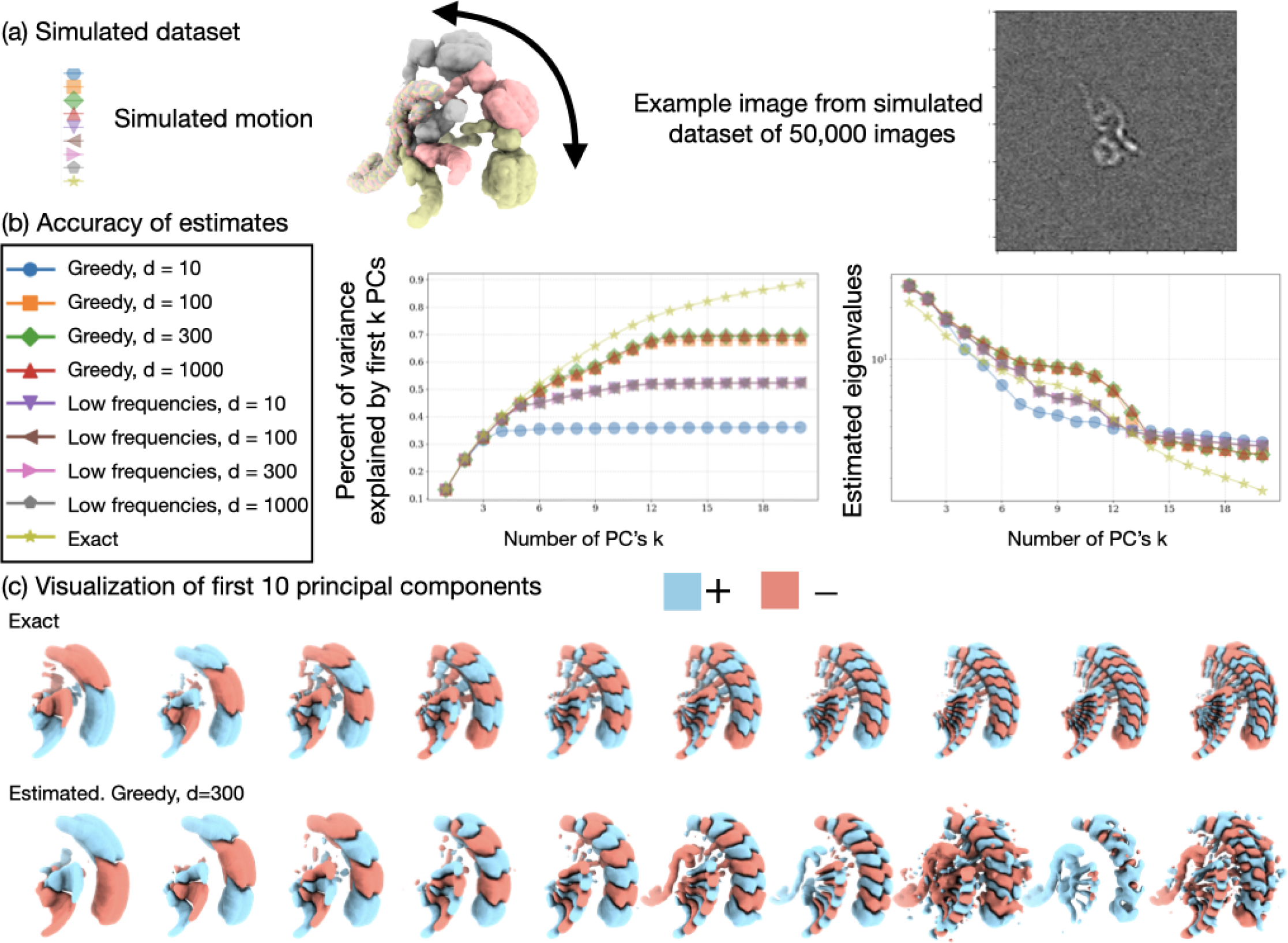
Accuracy of the proposed PCA method on a synthetic dataset. **(a)** Simulated dataset: a synthetic motion described in [65], and a synthetic dataset of 50,000 images. **(b)** Accuracy of principal components and estimated eigenvalues for different parameters. We compare the accuracy of two column sampling strategies: “Greedy” is the one described in appendix A.2, and “Low frequencies” is picking the columns corresponding to the lowest frequency voxels. We also show accuracy from computing a different number of columns *d* = 10, 100, 300, 1000 (before doubling using the Hermitian symmetry). The accuracy of the principal component is described using percent of the captured variance, computed as 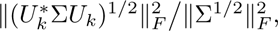 where *U*_*k*_ is the matrix containing the first *k* estimated principal components, Σ is the true covariance matrix and Σ^1/2^ is the matrix square-root, so that 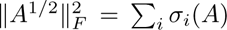 where *σ_i_*(*A*) are the singular values of *A*. Thus, the first 20 exact principal components capture 90% of the variance (which is the optimal choice), whereas the estimated using (Greedy, *d* = 300) captures around 70% of the captured variance. Here, the “Low frequency” sampling scheme achieves similar performance for all values of *d* (around 50% of captured variance), and the “Greedy” scheme improves with *d* until around *d* = 100 (around 70% of captured variance). The default value in RECOVAR is *d* = 300, which we observed empirically to be robust to a wide range of distributions. The scheme slightly overestimates the eigenvalues by a factor of around 1.2, possibly due to a misestimation of the mean and noise distribution which inflates the covariance estimate. Nevertheless, the decay of eigenvalues is correctly estimated until around the 12th eigenvalue where the noise dominates. **(c)** Visualization of the first ten exact and estimated principal components. The exact principal components are increasing in frequency, oscillating between positive and negative in the direction of motion. The estimated principal components increase in frequency but are increasingly noisy.

**Figure A.7:**
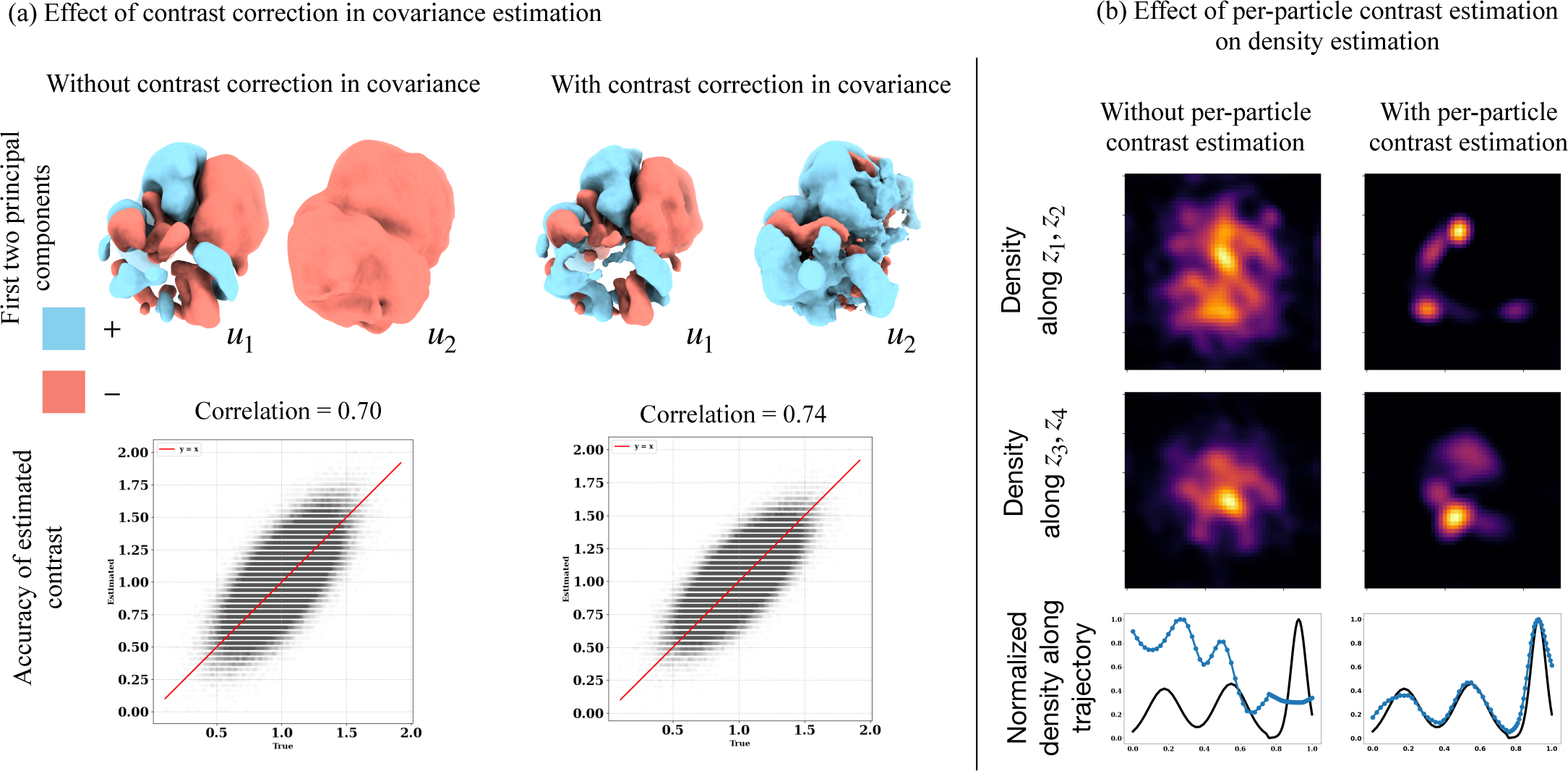
Illustration of the effect of contrast correction on the dataset in section 1.6. **(a)** Effect of contrast correction in covariance estimation as described in appendix A.4 on the computed principal components and per-particle correction. The second principal component estimated without contrast correction is strictly negative (color red) and reflects the contrast variation in the dataset. After contrast correction, the second principal component correctly encodes a motion. The contrast correction on the covariance matrix also improves the estimation accuracy of the per-image contrast estimation done in appendix A.5. **(b)** Effect of per-particle contrast estimation in appendix A.4 on density estimation. The conformational density is estimated with and without per-particle contrast estimation over 4 dimensions and is plotted over pairs of axes by integrating along the remaining axes. Also plotted is the estimated density along the top-down trajectory vs the ground truth density. (see fig. A.8 for more details). Without per-particle contrast estimation, the variation, in contrast, is embedded in latent space along with the heterogeneity, resulting in poor interpretability of latent space and inaccurate density estimates.

**Figure A.8:**
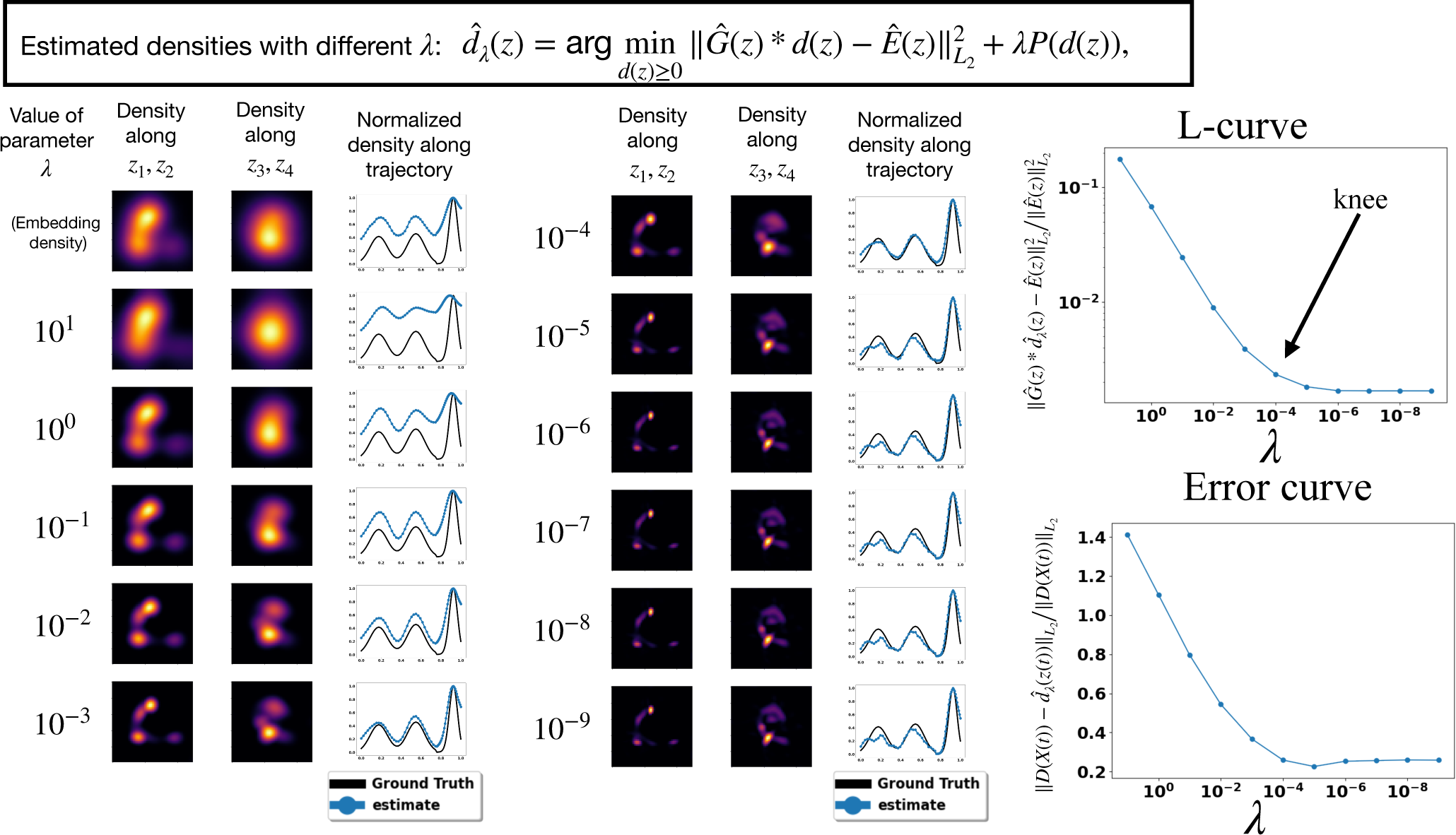
Illustration of density estimation with different regularization parameters on the synthetic dataset described in section 1.6. **(Left)** Embedding density and estimated densities with 11 different values of the parameter *λ*. The density is estimated over 4 dimensions and is plotted over pairs of axes by integrating along the remaining axes. Also plotted is the estimated density along the top-down trajectory vs the ground truth density. **(Right)** Computed L-curve and error curve between the true density and estimated along the true trajectory. The L-curve can be computed without knowledge of the ground truth as it only uses the residual. The L-curve criterion identifies the knee in the L-curve as the parameter choice for the inverse problem, here around *λ* = 10^*−*4^. This value is used to generate figures in fig. 2.

**Figure A.9:**
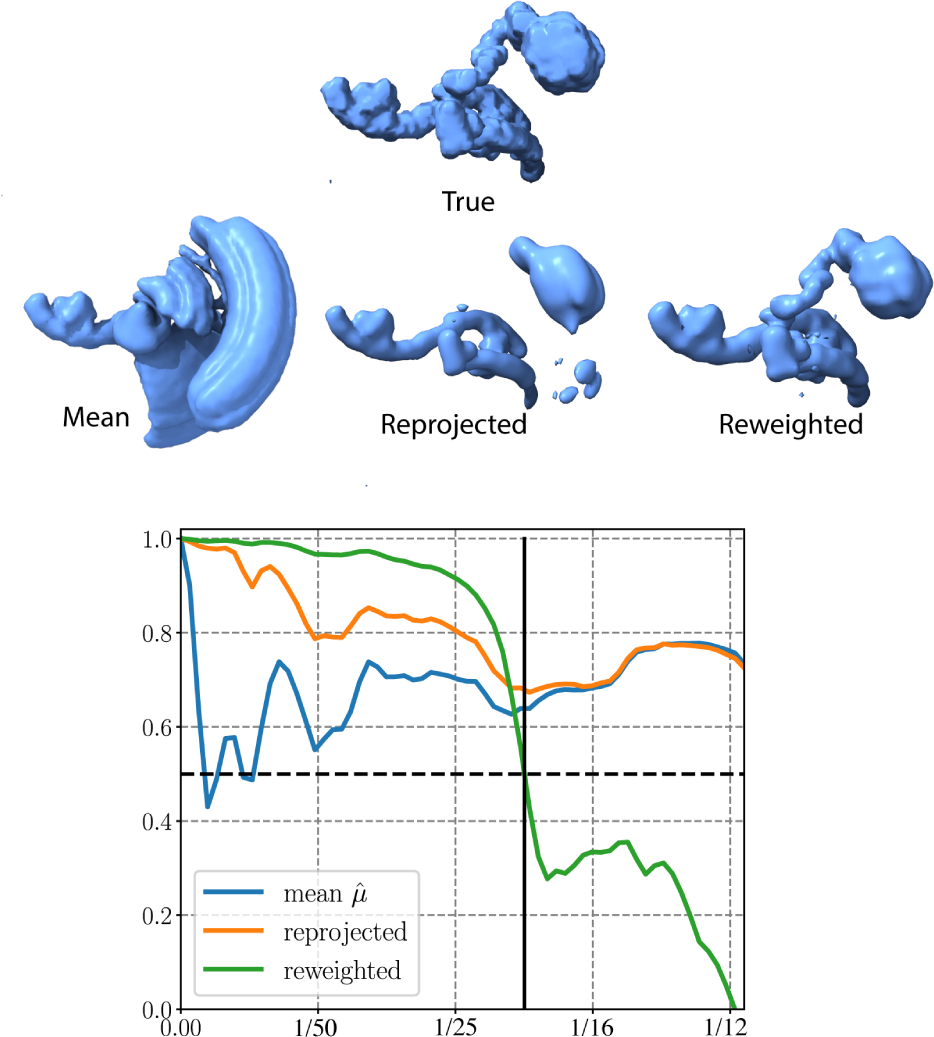
Illustration of the deceptive nature of FSC scores in heterogeneity analysis. **Top:** the true state, the estimated mean conformation, and reconstructed states using reprojection (“reprojected”) and fixed width kernel regression (“reweighting”) from the dataset detailed in [65] (’uniform’). **Bottom:** Fourier Shell Correlation curves, computed with a loose mask, between the true state and the reprojected reconstruction (orange), the fixed width kernel regression (green), and the estimated mean conformational state (blue).

**Figure A.10:**
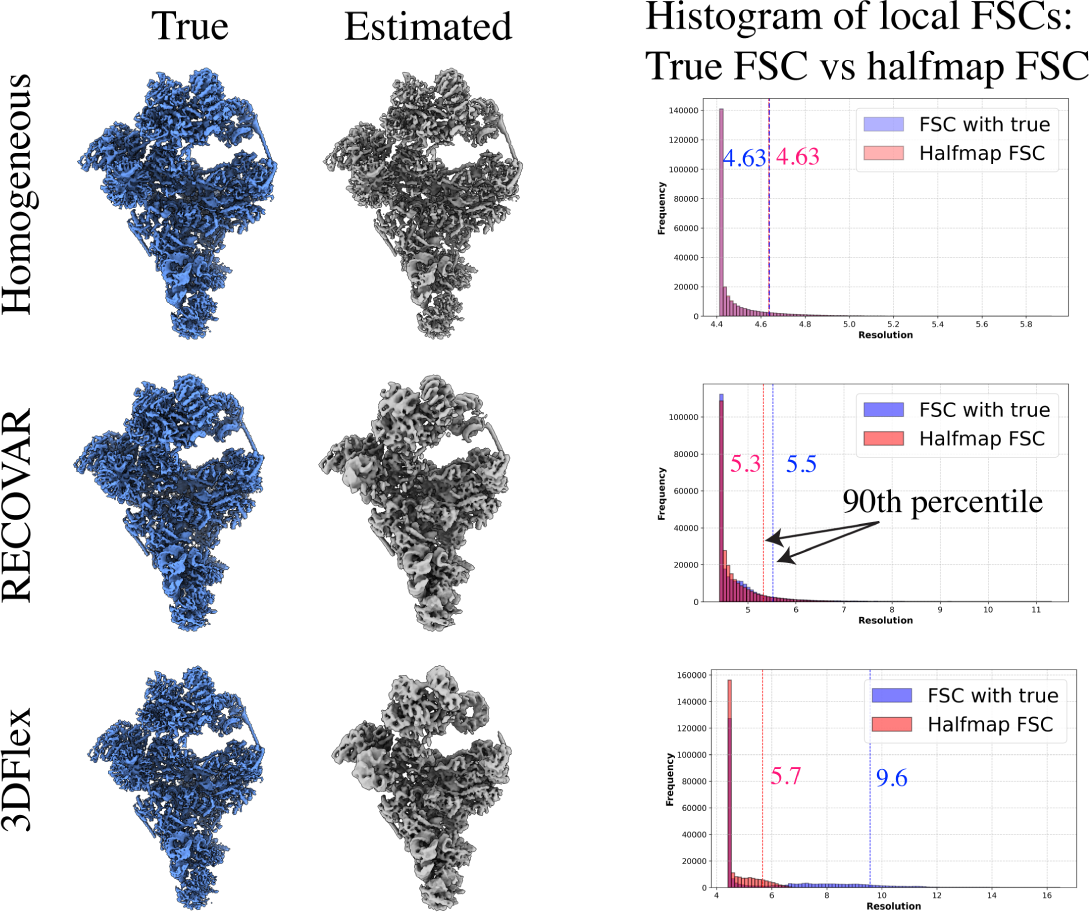
Illustration of the bias of the “gold-standard” FSC scores computed from halfmaps in heterogeneous reconstructions. **Top:** A homogeneous reconstruction of a state from a homogeneous synthetic dataset. The histogram of local FSC between the true state and the combined state (FSC threshold of 1*/*2) and the histogram of the FSC between halfmaps (FSC threshold of 1*/*7) overlap perfectly, as the halfmap FSC is an unbiased estimate of the true FSC. **Middle:** A RECOVAR reconstruction of the highest density state in the synthetic dataset in section 1.6. In this case, the histograms do not overlap as the halfmap FSC is a biased estimator of the true FSC and overestimates the resolution. **Bottom:** The 3Dflex “flexible reconstruction” of the whole synthetic dataset in section 1.6, compared to the most similar of the 1640 true volumes (as measured by the 90% percentile of local resolution score). The halfmap FSC of the flexible reconstruction vastly overestimates the true resolution of the flexible reconstruction.

**Figure A.11:**
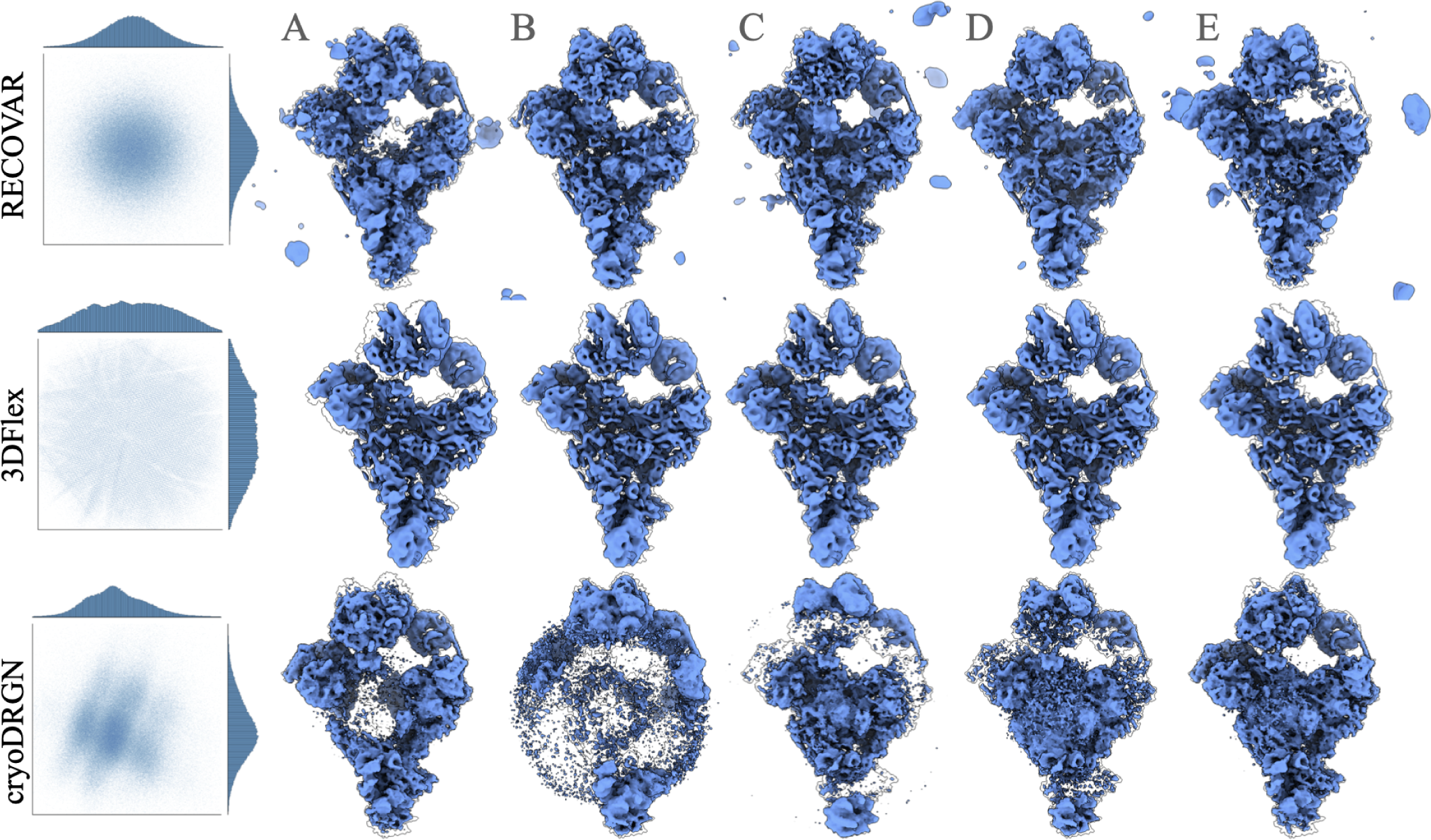
Robustness and hallucination of heterogeneity methods on a synthetic homogeneous dataset. Result of RECO-VAR, 3DFlex, and cryoDRGN on a similar dataset as in section 1.6 except without heterogeneity (all images are of the same conformation), and noise variance increased by 10. For all methods, we compute an embedding using default parameters. **(Left)** The first two PCs of the embedding are shown for each method. **(Right)** A motion generated by the default output by each method. For RECOVAR, it is the output of the “analyze” command, using the first path generated, for 3DFlex it is the motion generated along the first latent variable using the “3DFlex Generator” job, and for cryoDRGN, it is the motion along the first PC by the “analyze” command. The true volume is outlined with each state. All three methods show signs of hallucinated heterogeneity, RECOVAR appears to show extra densities around the volume, and some disappearing density (state E) could be incorrectly interpreted as compositional heterogeneity. 3DFlex hallucinates large (40− 50*Å*), high-resolution and physically-plausible motion. CryoDRGN shows variation which would likely be interpreted as imaging artifacts.

**Figure A.12:**
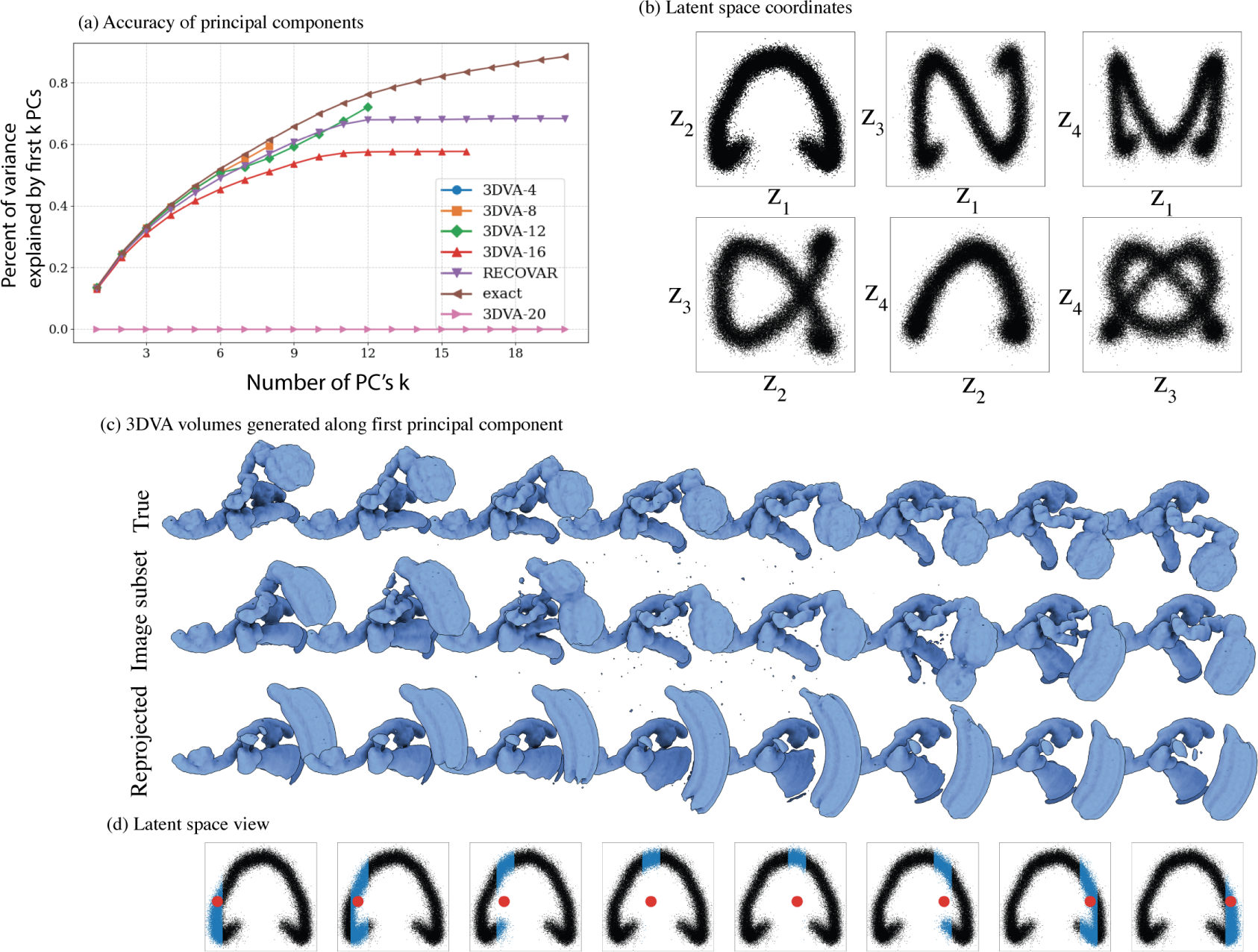
Result of 3DVA on a synthetic dataset with large motion(see [65], ‘uniform’) with a loose mask. **(a)** Accuracy of the computed principal components for different choices of dimension (3DVA-4,8,12,16,20) and for RECOVAR. The accuracy metric used is the percent of captured variance in the subspace. At 20 dimensions, the lack of regularization of 3DVA causes instability in the algorithm, and the program returns an error (depicted as 100% error). **(b)** The estimated latent coordinates using dimension 4. **(c)** True trajectory and reconstruction along the first principal component using an image subset averaging strategy (option “intermediate” in cryoSPARC) and reprojected (option “linear” in cryoSPARC). **(d)** Latent depiction of the motions recovered by 3DVA: in blue are the particles used in the subset averaging scheme, and in red is the latent coordinate used to reproject the first principal component. In either case, large artifacts are present due to the averaging of particles from distinct states in the case of subset averaging and the early truncation of the principal component in the reprojection case.

**Figure A.13:**
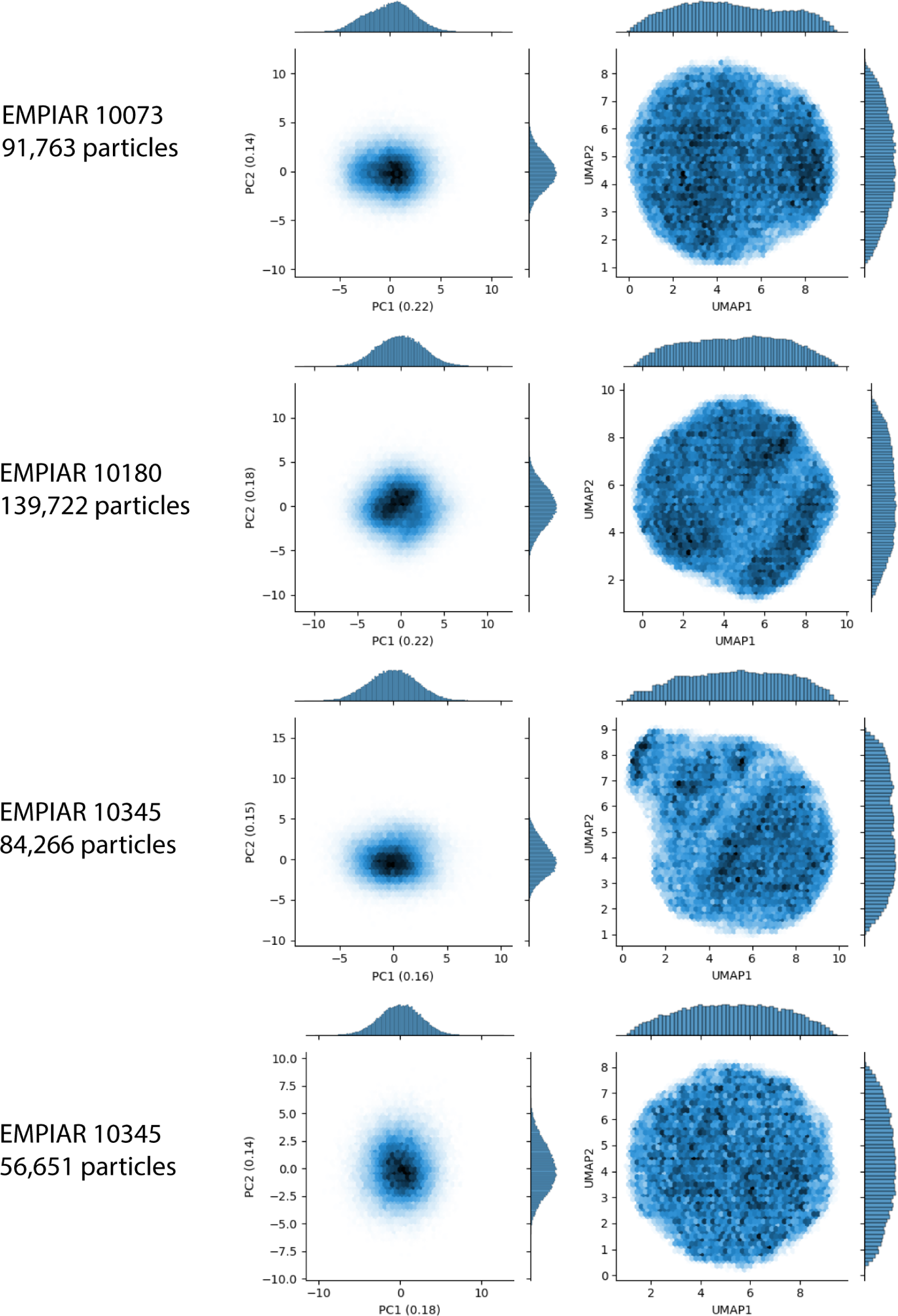
Embedding of cryoDRGN on the experimental datasets for comparison. Default parameters were used for all cases *β* = 1*/*8, 25 epochs.

## Code availability

The software is available at https://github.com/ma-gilles/recovar, and will be incorporated in the ASPIRE software system [62].

## Data availability

Data for synthetic experiment in section 1.6 will be deposited on https://zenodo.org. Experimental datasets are deposited in EMPIAR with entry ID 10076, 10180, 10073 and 10345. Filtered indices, and poses the datasets 10076 and 10180 are deposited in https://zenodo.org/records/10030109 and for 10073 and 10345 will be deposited on zenodo https://zenodo.org.

## Acknowledgments

The authors thank Eric Verbeke, Joakim Andén, Roy Lederman, Edgar Dobriban, William Leeb, Bogdan Toader, Willem Diepeveen, Ellen Zhong, Pilar Cossio, Luke Evans, Lars Dingeldein and Erik Thiede for helpful discussions. Large part of our implementation is adapted from the software suites RELION and cryoDRGN, and the authors thank the developers for making the software and code accessible. The authors also thank the developers of cryoSPARC, EMDA, and ChimeraX for making their software freely available, all of which were used to generate the results presented here. The density estimation in section 1.4 was initially developed to address the inaugural Flatiron Institute cryo-EM heterogeneity community challenge, and the authors gratefully acknowledge the organizers for inspiring this line of work.

The authors are supported in part by AFOSR under Grant FA9550-23-1-0249, in part by Simons Foundation Math+X Investigator Award, in part by NSF under Grant DMS-2009753, and in part by NIH/NIGMS under Grant R01GM136780-01. Part of this research was performed while the first author was visiting the Institute for Pure and Applied Mathematics (IPAM), which is supported by the National Science Foundation (Grant No. DMS- 1925919).

## Competing interests

The authors declare no competing interests.

## Footnotes

1 The spatial domain variance map is displayed in fig. 1 only for illustration, the regularization is computed between columns of the covariance matrix in the Fourier domain, see appendix A.3.

2 Timings were performed on an NVIDIA A100 80GB GPU.

3 The embedding is the set of pairs 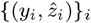, which assigns each image *y*_*i*_ to its mean label 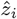.

4 This point is also made in the paper titled “Principal component analysis is limited to low-resolution analysis in cryoEM” [54] which argues the reprojection scheme used with 2-3 principal components produces low-resolution volumes. In our language, it is not PCA itself that is low-resolution, but rather the reprojection scheme.

5 We note that it is the density in atomic coordinates that is related to the Boltzmann distribution. Cryo-EM methods reconstruct volumes, not atomic coordinates and these two densities do not exactly coincide, though they are related. We ignore this issue here and focus on computing the density of volumes.

6 For cryoDRGN we test the parameters *β* = 1*/*8, **1** and EPOCHS = 12, 25, **50**, and 3DFlex test the parameters dim(*z*) = 2, **3** and rigidity = 0.2, **2**, 20. The best parameters, used for benchmarking, are highlighted in bold.

7 For RECOVAR, states are embedded using the PCA projection map. For cryoDRGN and 3DFlex, states are embedded in latent space by the median label of images of that state, and the density is computed from a kernel density estimator with the Silverman rule. The true density *D*(*X*(*t*)) along the trajectory is parametrized by 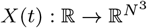 by enforcing 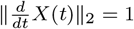.

8 Similarly, the cryoDRGN embedding shows clusters in the unfiltered stack but none in the filtered stack, see fig. A.13.

9 We denote grid frequencies with superscript *ξ*^*k*^ and off-the grid frequencies sampled by images with subscript *ξ*_*i,j*_ to make the distinction clearer.

10 To our knowledge, this link with kernel regression and RELION was first noted in [28]

11 We compute the SVD in the spatial domain using a randomized SVD [30] and truncate the rank of the SVD to rank 200 if *d >* 200 by default.

12 While some of these methods could be used directly, we find empirically that they work poorly under the highly anisotropic noise distribution in the entries of 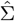, which is why we devise the projected covariance scheme above.

13 Notably, this iteration does not necessitate iterating through the data. Iterating reduces the variance of the final estimator.

14 This assumption is often violated, as the principal components are often not exactly orthogonal to the mean. However, we have found empirically that it improves the contrast estimation to make this correction. The same assumption is made in [55].

15 In some cases, it is preferable to consider the (unregularized) maximum-likelihood label (e.g., in appendix A.7). In that case, we take Γ^*−*1^ = 0 in eq. (22)

16 We do not use images for which 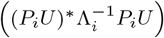 is rank deficient or highly ill-conditioned in what follows.

17 Our implementation uses scikit-fmm, see https://github.com/scikit-fmm.

18 In both PPCA and 3DVA, *U* is not constrained to be orthogonal.

19 Under these assumptions, the unregularized covariance approach is equivalent to PCA

20 In [57], it is shown that EM applied to the PPCA problem is globally convergent if *P*_*i*_ ≠ *I*, but the proof does not treat the case relevant to cryo-EM where *P*_*i*_ = *I*.

21 The total complexity of RECOVAR is 𝒪(*nN*^2^*d* + *N*^3^*d*^2^ + *nd*^4^ + *d*^6^), and of PPCA and 3DVA is 𝒪(*nN*^2^*d*^2^s + *N*^3^*d*^2^s). They also have different memory complexity. Without counting the dataset of size 𝒪(*nN*^2^), their memory complexity is 𝒪(*nd* + *N*^3^*d*^2^) for 3DVA, 𝒪(*nd*^2^ + *N*^3^*d*^2^) for PPCA, 𝒪(*nd*^2^ + *N*^3^*d*) for RECOVAR. The difference in storage is often small compared to the size of the image dataset.

## Notes

https://github.com/ma-gilles/recovar

### Competing Interest Statement

The authors have declared no competing interest.

### Summary of Updates

Fixed error in complexity analysis, which incorrectled stated the complexity of RECOVAR, PPCA, 3DVA.

